# Accurate Bayesian segmentation of thalamic nuclei using diffusion MRI and an improved histological atlas

**DOI:** 10.1101/2022.09.28.508731

**Authors:** Henry F. J. Tregidgo, Sonja Soskic, Juri Althonayan, Chiara Maffei, Koen Van Leemput, Polina Golland, Anastasia Yendiki, Daniel C. Alexander, Martina Bocchetta, Jonathan D. Rohrer, Juan Eugenio Iglesias, the Alzheimer’s Disease Neuroimaging Initiative

## Abstract

The human thalamus is a highly connected brain structure, which is key for the control of numerous functions and is involved in several neurological disorders. Recently, neuroimaging studies have increasingly focused on the volume and connectivity of the specific nuclei comprising this structure, rather than looking at the thalamus as a whole. However, accurate identification of cytoarchitectonically designed histological nuclei on standard *in vivo* structural MRI is hampered by the lack of image contrast that can be used to distinguish nuclei from each other and from surrounding white matter tracts. While diffusion MRI may offer such contrast, it has lower resolution and lacks some boundaries visible in structural imaging. In this work, we present a Bayesian segmentation algorithm for the thalamus. This algorithm combines prior information from a probabilistic atlas with likelihood models for both structural and diffusion MRI, allowing label boundaries to be informed by both modalities. We present an improved probabilistic atlas, incorporating 26 thalamic nuclei identified from histology and 45 white matter tracts identified in ultra-high gradient strength diffusion imaging. We present a family of likelihood models for diffusion tensor imaging, ensuring compatibility with the vast majority of neuroimaging datasets that include diffusion MRI data. The use of these diffusion likelihood models greatly improves identification of nuclei versus segmentation based solely on structural MRI. Dice comparison of 5 manually identifiable groups of nuclei to ground truth segmentations show improvements of up to 10 percentage points. Additionally, our chosen model shows a high degree of reliability, with median test-retest Dice scores above 0.85 for four out of five nuclei groups, whilst also offering improved detection of differential thalamic involvement in Alzheimer’s disease (AUROC 83.36%). The probabilistic atlas and segmentation tool will be made publicly available as part of the neuroimaging package FreeSurfer.

## 1. Introduction

The thalamus has traditionally been considered a relay station for information in the brain, with extensive connections to both cortical and subcortical structures (Schmahmann, 2003; Behrens et al., 2003). As such, it integrates information processing between cortical regions (Sherman, 2007, 2016; Hwang et al., 2017) and is associated with a wide range of functions including cognition, memory, sensory and motor functions, regulation of consciousness and spoken language among others (Sherman and Guillery, 2001; Schmahmann, 2003; Fama and Sullivan, 2015). Additionally, neurodegenerative pathological processes in the thalamus have been associated with Alzheimer’s disease (*AD*) (de Jong et al., 2008; Zarei et al., 2010), frontotemporal dementia (Bocchetta et al., 2018; McKenna et al., 2022), Huntington’s disease (Aron et al., 2003; Kassubek et al., 2005) and multiple sclerosis (Minagar et al., 2013; Planche et al., 2019).

With such wide established connections and functions, the thalamus is a frequent target in MRI-based neuroimaging studies and a focus for research in relation to both healthy and disordered brain function. This creates a need for reliable identification of thalamic borders. Therefore, the thalamus is defined by several structural MRI (*sMRI*) segmentation methods, including multi-atlas segmentation (Heckemann et al., 2006), Bayesian segmentation (Puonti et al., 2016) and convolutional neural networks (*CNNs*) (Wachinger et al., 2018; Billot et al., 2020; Henschel et al., 2020). Additionally, the thalamus has been included in popular image processing packages, including FreeSurfer’s (Fischl, 2012) recon-all stream, which uses a probabilistic atlas of anatomy and MRI intensity (Fischl et al., 2002), and the FMRIB Software Library (*FSL*) (Smith et al., 2004), which includes a model of shape and appearance in its implementation (FIRST) (Patenaude et al., 2011).

The methods above segment the thalamus as a single label, however in reality it is a complex and heterogeneous structure. It is composed of 14 major nuclei, which may be split further into 50 subnuclei depending on the level of detail in the classification and agreement on neuroanatomical definition. There are multiple such definitions with varying numbers of subnuclei (Morel, 2007; Jones, 2012; Mai and Majtanik, 2019). These nuclei have distinct patterns of connections with other brain regions and subserve different functions, including associative, sensory, motor, cognitive and limbic (Schmahmann, 2003). For example, the ventral lateral posterior nucleus is involved in motor function through connections with the cerebellum and the motor cortex, while the mediodorsal nucleus has connections with the prefrontal cortex and plays a role in cognitive and emotional processes (Mai and Forutan, 2012; Schmahmann, 2003). In addition, neuropathological studies have demonstrated preferential involvement of certain thalamic nuclei in several conditions, such as the caudal intralaminar nuclei in Parkinson’s disease (Henderson et al., 2000), the anterior nuclei in AD (Braak and Braak, 1991a,b), and the pulvinar in the *C9orf72* genetic subtype of frontotemporal dementia (Vatsavayai et al., 2016). These studies provide strong motivation for the design of automated segmentation algorithms that accurately define thalamic nuclei *in vivo*, enabling identification of reliable and precise biomarkers.

Different approaches have been used to segment thalamic nuclei. There are segmentation strategies that attempt to directly register histology derived labels to MRI. For instance, manually labelled histology can be used to generate a reference space atlas that may then be applied to *in vivo* MRI through registration-based segmentation (Krauth et al., 2010; Jakab et al., 2012; Sadikot et al., 2011). However, such approaches are limited by the difficulty in registering MR images with different contrasts. Other techniques define their label scheme based on information derived from the imaging data to be segmented. For example, diffusion MRI (*dMRI*) has been used to define thalamic regions by clustering voxels based on diffusion tensor imaging (*DTI*) indices (Mang et al., 2012) and orientation distribution functions (Battistella et al., 2017; Semedo et al., 2018). Other studies have divided the thalamus into regions based on their cortical connectivity, either through resting-state functional MRI time course correlations (Zhang et al., 2008) or dMRI tractography (Behrens et al., 2003; Johansen-Berg et al., 2005). However, exactly how thalamic regions defined by functional MRI relate to neurobiology is not fully understood (Eickhoff et al., 2015) and there is some indication that tractography-based segmentations are insensitive to the internal structure of the thalamus (Clayden et al., 2019).

The development of advanced MRI acquisitions has also allowed for atlases to be defined from manual segmentation of *in vivo* imaging directly, due to improved resolution and contrast. For example, guided by histological atlases, it has been possible to manually identify nuclei on advanced Smri acquired at 7T (Tourdias et al., 2014; Liu et al., 2020) and on dMRI through short-track track density imaging (Basile et al., 2021). In particular, segmentations of 7T white-matter-nulled imaging have been used to generate both multi-atlas segmentation (“THOMAS” Su et al. 2019) and CNN (Umapathy et al., 2021) segmentation algorithms. However, these segmentations do not have the full level of detail present in histological atlases and performance is impacted by changes in acquired contrast, due to domain gap effects for CNNs and poorer registration in multi-atlas segmentation.

Aiming to provide detailed segmentations of thalamic nuclei that is robust to changes in MRI acquisition and contrast, we previously constructed a probabilistic atlas of the thalamus and surrounding tissue from manually labelled histology (Iglesias et al., 2018). We then combined this atlas with Bayesian inference methods (Wells et al., 1996; Van Leemput et al., 1999; Ashburner and Friston, 2005; Pohl et al., 2006) to allow segmentation of 25 bilateral histological labels from sMRI. This approach had the advantage that the intensity model of each label was learned from the target image, allowing the resulting labels to remain stable across acquisition contrasts. However, these segmentations can be less accurate in areas where sMRI shows poor contrast. For example, Fig. 1 shows that our previous method can accurately follow the boundary between groups of medial and lateral nuclei, but the lack of contrast between lateral nuclei and white matter can lead to oversegmentation into the internal capsule.

**Figure 1:**
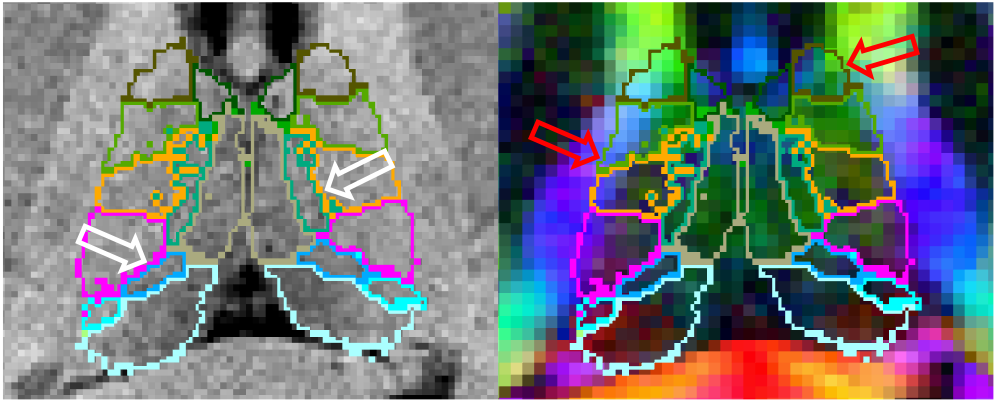
Thalamic segmentation of a T1-weighted structural MRI overlaid on the co-registered T1-weighted image (left) and a co-registered directionally encoded colour FA image (right). High contrast between medial and lateral thalamic regions on structural imaging improves the accuracy of these boundaries (white arrows). However, low contrast between the lateral thalamus and white matter causes over-segmentation into the internal capsule, which can easily be discerned in the colour FA image (red arrows).

The availability of complementary information from dMRI sequences provides a possible avenue for minimising such segmentation errors. An increasing number of large multi-site neuroimaging studies, including the Human Connectome Project (*HCP*) (Van Essen et al., 2013), the Alzheimer’s Disease Neuroimaging Initiative (ADNI) (Jack Jr. et al., 2008), and the GENetic Frontotemporal dementia Initiative (GENFI) (Rohrer et al., 2015) are acquiring both structural and diffusion MRI. Additionally, use of DTI combined with registration-based segmentation has been proposed for segmentation of the whole thalamus in subjects where T1-weighted MRI contrast is very low (Al-Saady et al., 2022). As can be seen in Fig. 1, dMRI shows good contrast between the thalamus and the adjacent white matter, while structural MRI provides better contrast between the medial nuclei and cerebrospinal fluid (*CSF*) as well as higher resolution. Therefore, we look towards creating joint models of structural and diffusion MRI, incorporating likelihood models of DTI such as those used in the modelling of white matter fibres (Jian and Vemuri, 2007).

We present an extension of our structural Bayesian inference segmentation algorithm to incorporate dMRI. We focus on DTI due to the ease of fitting tensors to diffusion-weighted images, even from legacy data or in studies with short acquisitions. We explore our recently proposed diffusion likelihood model, combining the Dimroth-Scheidegger-Watson (*DSW*) and Beta distributions (Iglesias et al., 2019). We compare this model to both the Wishart distribution, from fibre modelling literature (Jian and Vemuri, 2007), and the log-Gaussian distribution, influenced by tensor interpolation methods (Arsigny et al., 2006). Additionally, we build on our previous histological atlas of the thalamus by adding 45 labels for white matter tracts passing adjacent to the thalamus, allowing the DTI likelihood models to capture the varying directionality of fibers in white matter without becoming sensitive to non-white-matter tissue. The resulting segmentation method allows constraints to be imposed independently on both the structural and diffusion modelling by including separate shared parameter models, enforcing reflective symmetry, incorporating prior distributions on likelihood parameters, and re-weighting likelihood terms to account for the lower resolution of DTI.

This paper is structured as follows. In Section 2 we outline our joint segmentation method. This includes explanations of: the general Bayesian inference model; the model fitting and segmentation process; the three likelihood models; the atlas and its construction; and general implementation details. In Section 3 we evaluate our joint segmentation method on both high and low resolution data. This evaluation includes: model optimisation and evaluation on a population template constructed from both T1-weighted MP-RAGE and DTI images; evaluation of the optimised models on subjects from HCP, providing comparison to manual ground truth and test-retest reliability; and test-retest and indirect evaluation on conventional quality data. Section 4 concludes the paper.

## 2. Bayesian segmentation of brain MRI

### 2.1. Probabilistic model and Bayesian inference

Here we outline the theory and implementation of our Bayesian segmentation algorithm. As in existing Bayesian segmentation literature (Van Leemput et al., 1999; Zhang et al., 2001; Ashburner and Friston, 2005; Iglesias et al., 2015; Puonti et al., 2016), our strategy relies on modelling the voxel-wise data as observable random variables. These follow a different distribution for each label class in a supplied deformable probabilistic atlas of the volume encompassing the thalamus (Van Leemput, 2009; Iglesias et al., 2018). Both the voxel-data distributions and deformation of the atlas are parameterised by hidden random variables dependent on the subject and image acquisition. Estimating these hidden random variables allows us to generate a voxel-wise probability of membership in each label class (Van Leemput et al., 1999; Ashburner and Friston, 2005). In the Bayesian approach, this is used to construct the posterior probability of a labelling (or segmentation) given paired sMRI and dMRI data.

For the purposes of this method we assume that both the sMRI and dMRI have been registered and resampled to the same grid comprised of voxels indexed by *υ* ∈ {1, …, *V* }. We denote the labelling of these voxels by *L* = [*l*_1_, …, *l*_*V*_], with *l*_*υ*_ ∈ {1, …, *C*} – where *C* is the number of label classes in our model. Similarly, we construct a matrix *S* = [*s*_1_, …, *s*_*V*_] holding vectors of sMRI voxel data, *s*_*υ*_, and matrix ***d*** = [***d***_1_, …, ***d***_*V*_] to hold the dMRI voxel data, ***d***_*υ*_. We explore different representations of ***d***_*υ*_ in later sections.

Using this notation and applying Bayes’ rule, the posterior probability of a specific labelling for a pair of sMRI and dMRI scans of a subject is:

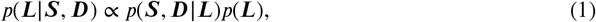

and the labelling that maximises Eq. (1) is known as the maximum a posteriori (*MAP*) estimate for the segmentation. To obtain this MAP estimate we need both the *likelihood* distribution, *p*(*S, D*|*L*), of our imaging data given a segmentation, and a *prior* distribution, *p*(*L*), generated from prior anatomical knowledge of the thalamus and its surroundings. As these can be used to generate random scans by sampling first from the prior then from the likelihood, segmentation can be thought of as fitting a generative probabilistic forward model to our data and “inverting” it to obtain the labelling.

To make the problem in Eq. (1) tractable, we assume: *i)* that both the likelihood and prior factorise over voxels and *ii)* that the sMRI and dMRI are independent of each other given the labels. The exact graphical model of our framework is shown in Fig. 2. At the top of this model we define the prior distribution on the labels, beginning with a probabilistic atlas *A*.This atlas is constructed within a reference brain space, meaning it is likely to match the topology of any segmentation subject, but will require deformation to match accurately. The atlas *A* provides, at each spatial location, the prior probability of observing each neuroanatomical label class. We define *A* on a deformable tetrahedral mesh, where each vertex has an associated vector of class probabilities, and barycentric interpolation can be used to obtain probabilities at non-vertex locations (Van Leemput, 2009). We define a set of parameters, *θ*^*a*^, that move the mesh nodes to deform the atlas into the space of the target MRI voxel grid, accommodating the anatomical variability across subjects. These parameters are themselves a sample from a distribution that is regularised by setting the stiffness *γ*^*a*^, preventing folding of the atlas mesh and preserving topology. We then assume that our labelling *L* is sampled from the categorical distribution over classes defined by the deformed atlas, with each voxel location sampled independently allowing factorisation.

**Figure 2:**
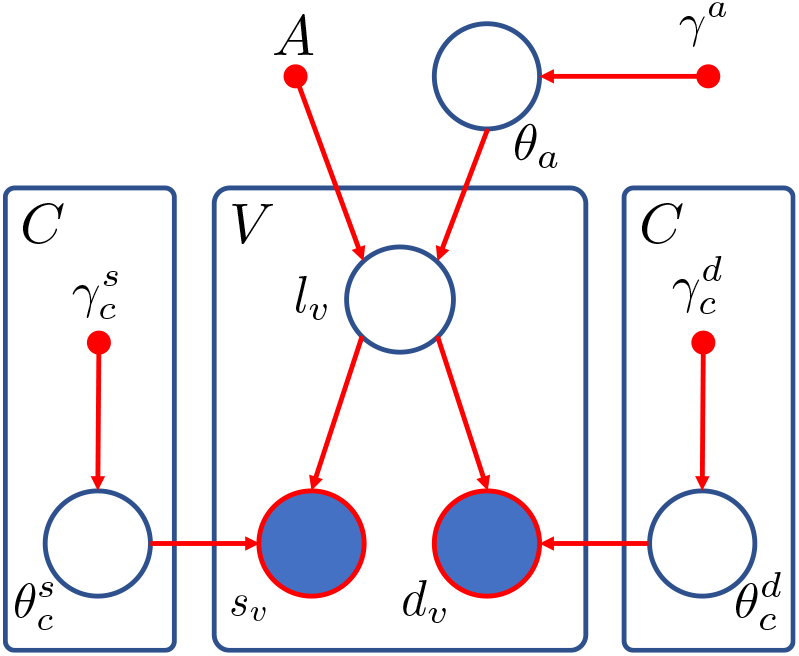
Graphical model of the proposed framework. Larger circles represent random variables with open circles for the hidden variables (*θ, l*), and shaded circles for the observed variables (*s,d*). Smaller solid circles are deterministic parameters such as the atlas (*A*) and encoded prior information (*γ*). Rectangles indicate replication across voxels (*V*) or classes (*C*).

Given *L* we can define the likelihood model for our observed data. We assume that the sMRI and dMRI are conditionally independent from each other and across voxels given the labelling, with *s*_*υ*_ and ***d***_*υ*_ modelled as samples from separate distributions parameterised by 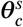 and 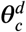 respectively. These hidden parameters are dependent on the corresponding label *l*_*υ*_ = *c*. Any prior knowledge on these parameters is encoded in prior distributions controlled by hyperparameters 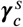 and 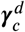.

Under these assumptions we can define the full joint probability density function (*PDF*) for Fig. 2 as

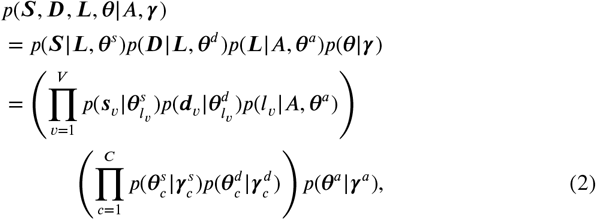

where 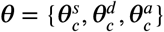 and 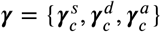.

With the model described by Fig. 2 and Eq. (2) we can formulate the MAP estimate for our segmentation as

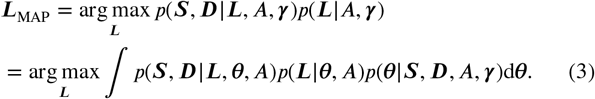

However, integrating the joint PDF over the full space of possible parameters *θ* is intractable. For this reason we make the standard assumption that the posterior distribution of the hidden parameters is heavily peaked around the mode, 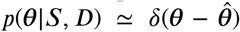. In this way, we can segment our images by applying Bayes’ rule to Eq. (2) and marginalising over the hidden labelling *L* to obtain these optimal hidden parameters (so called “point estimates”):

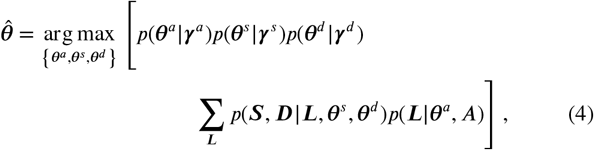

and then optimising *L* to obtain the MAP estimate

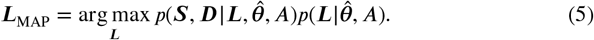

### 2.2. Parameter estimation and segmentation

The first step in the segmentation process is to estimate the optimal hidden parameters 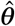 from Eq. (4). We begin by formulating the likelihood PDFs for both sMRI and dMRI as mixture models such that each label class is described by mixtures of *G* structural component distributions and *W* diffusion component distributions, giving

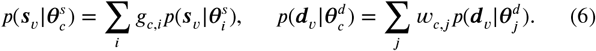

Here, *g*_*c,i*_ ≥ 0 and *w*_*c,j*_ ≥ 0 are mixture weights in the model of label class *c* indicating the contribution of the *i*-th sMRI and *j*-th dMRI component’s distribution to the appearance of the class in the respective modality. These distributions are parameterised by 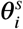 and 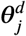, respectively, with *i* ∈ 1, …, *G* and *j* ∈ 1, …, *W*. In both cases the sum over the component weights for a given class must be equal to one, ∑_*i*_ *g*_*c,i*_ = 1 and ∑_*j*_ *w*_*c,j*_ = 1. This mixture model formulation provides a high degree of flexibility, allowing us to specify *a priori* which label classes may be modelled jointly by constraining specific weights to 0 or 1 while others are allowed to vary.

Combining Eq. (6) with Eqs. (2) and (4) and taking logarithms we can then obtain an objective function to be optimised with respect to the distribution parameters,

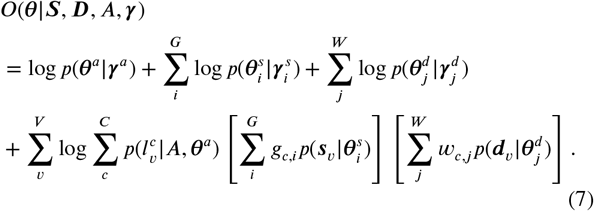

To optimise Eq. (7) we adapt the approach proposed by Puonti et al. (2016). In this approach the atlas deformation parameters and likelihood parameters are optimised iteratively in a coordinate ascent scheme, with each being optimised while the other is fixed. The optimisation of the *θ*^*a*^ is performed using a standard conjugate gradient operator with the deformation prior *p*(*θ*^*a*^|*γ*^*a*^) taking the form of the penalty term defined by Ashburner et al. (2000). The likelihood parameters *θ*^*s*^ and *θ*^***d***^ are then optimised using a Generalised Expectation Maximisation (*GEM*) algorithm (Dempster et al., 1977; Van Leemput et al., 1999), iterating between expectation (*E*) and Maximisation (*M*) steps.

#### E step

In the E step, we build a lower bound *Q*(*θ*) for the objective function in Eq. (7) using Jensen’s inequality:

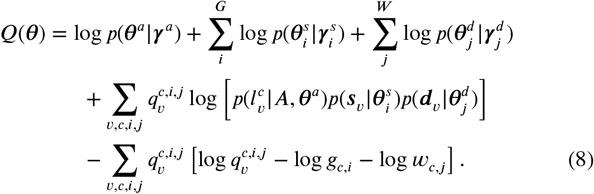

Here 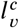 indicates the event that the voxel label *l*_*υ*_ = *c* and 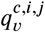 is a soft segmentation at the current parameter estimates indicating the combination of class *c*, sMRI distribution *i* and dMRI distribution *j*:

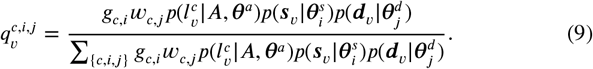

#### M step

In the generalised M step we attempt to increase the bound *Q*(*θ*) in Eq. (8). We note that the two sets of distribution parameters 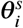 and 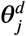 can be optimised individually, as they make independent contributions to the bound:

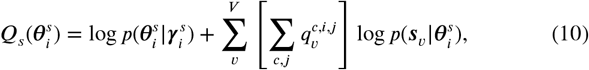

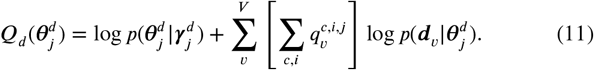

These contributions can then be optimised using either closed form solutions or numerical methods, depending on the distribution used as we will describe in Section 2.3. Finally we can calculate the new optimal weightings as

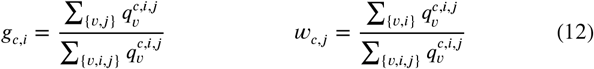

#### Segmentation

The mesh deformation and likelihood parameter optimisation steps are repeated alternately until the objective function in Eq. (7) has converged. At this point, we note that the formulation of the posterior factorises over voxels and the posterior probability of each class may be found by summing over the soft segmentations 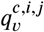. Hence the final MAP estimate segmentation is given by

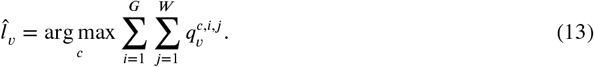

### 2.3. Likelihoods

So far, we have outlined the Bayesian framework and segmentation process without specifying the likelihood models used for both sets of MRI data. The steps outlined above are not affected by the choice of distributions used. Here we provide an overview of the distributions used to model the sMRI and dMRI data, including the likelihood term and, where applicable, the prior over its parameters. Detailed equations for the calculation of PDF values as well as the optimisation of model parameters, *θ*, may be found in Section S.1 of the supplement.

#### 2.3.1. Structural MRI model

To model the sMRI intensities, we follow the Bayesian brain MR segmentation literature and use a mixture of Gaussian intensity distributions (Ashburner and Friston, 2005; Zhang et al., 2001; Van Leemput et al., 1999). In this model the intensity values for each structural modality are held in the vector *s*_*υ*_ and the model parameters 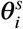 are the mean and covariance, {***μ***_*i*_, Σ_*i*_}, of the structural mixture component *i*. We choose to use the natural conjugate prior, the Normal-Inverse-Wishart distribution, on these Gaussian parameters. The likelihood and prior distributions are therefore

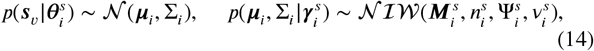

where 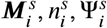 and 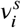 encode any prior knowledge we may have on the structural distribution. Formulations for the structural PDFs and closed form solutions to the parameter M step parameter optimisations can be found in Section S.1.1 of the supplement.

#### 2.3.2. Diffusion MRI models

To model the dMRI data, we consider distributions over tensors estimated with DTI. Even though higher-order models can be used with modern dMRI acquisitions, using DTI models ensures that our method is compatible with virtually every dMRI dataset, including huge amounts of legacy data. In this work, we compare two competing models, based on the Wishart and Gaussian distributions, to our previously-proposed DSW-beta distribution (Iglesias et al., 2019).

##### Wishart

Following existing white matter fibre modelling literature, we look to the Wishart distribution (Jian and Vemuri, 2007). DTI produces at each voxel a covariance matrix describing the displacements of water molecules in the voxel. Therefore, the natural conjugate prior for these tensors is an Inverse-Wishart distribution. We use this in combination with a Gamma distribution on the degrees of freedom parameter (Görür and Rasmussen, 2010), with the effect of lowering the degrees of freedom and increasing the breadth of the resulting Wishart distributions. In this model, we define ***d***_*υ*_ as the inverse of the diffusion tensor *T*_*υ*_. We then use the Wishart and Gamma distributions to model ***d***_*υ*_ and 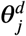:

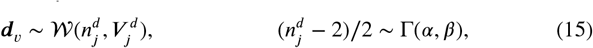

where *α* and *β* are set to 0.5 and 1.5 respectively to provide a non-informative prior. Formulations for the Wishart PDFs and the optimisation problem in the M step can be found in Section S.1.2 of the supplement.

##### Log-Gaussian

This model is motivated by literature on the interpolation of DTI volumes. Direct interpolation of DTI can lead to swelling of the ellipsoids representing the diffusion tensors, but interpolating in the log domain reduces this effect (Arsigny et al., 2006; Dryden et al., 2009). For this reason, and noting that the DTI tensors, *T*_*υ*_, are symmetric with only six independent variables, we define ***d***_*υ*_ as a vector

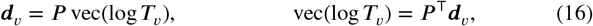

where *P* is a constant 6 × 9 matrix (values listed in supplement) designed with the constraint that

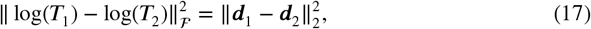

and therefore interpolation of the vectors ***d***_*υ*_ is equivalent to interpolation of the tensors in the log domain. In this formulation the natural distribution to choose based on the distance metric in Eq. (17) is a Gaussian distribution with a scalar variance

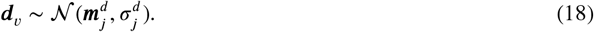

We then define uniform priors on both 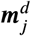 and 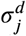 due to the difficulty in informing these parameters a priori. Formulations for the log-Gaussian PDFs and the optimisation problem in the M step can be found in Section S.1.3 of the supplement.

##### DSW-beta

This model is a custom distribution proposed in our prior work (Iglesias et al., 2019) motivated by a desire to lower the dimensionality of ***d***_*υ*_, leading to a reduction in extreme values of the likelihood that may overwhelm the prior. Here only the fractional anisotropy (*FA*), *f*_*υ*_, and the principal eigenvector, ***ϕ***_*υ*_, of the tensor *T*_*υ*_ are modelled so that ***d***_*υ*_ = {*f*_*υ*_, ***ϕ***_*υ*_}. In this approach, we use the two parameter Beta distribution to model the FA as it is able to model both the location and dispersion of signals in the range [0, 1]. We then use the DSW distribution to model ***ϕ***_*υ*_.

This DSW distribution has the advantages of symmetry and simplicity. As DSW is antipodally symmetric, it accommodates the directional invariance of dMRI (Zhang et al., 2012). It is also rotationally symmetric around a mean direction and its opposite {***ψ***, −***ψ*** : ‖***ψ***‖ = 1}, with a dispersion around the mean parameterized by a concentration *κ* (Mardia et al., 2000). This *κ* allows us to incorporate the higher directional dispersion in voxels with lower FA by multiplying the component specific concentration by the voxel FA to give an effective concentration for each voxel. The likelihood distribution in this formulation of the dMRI is therefore a joint DSW-beta distribution

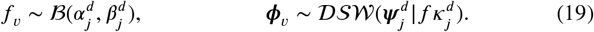

Formulations for the DSW-beta PDFs and M step can be found in Section S.1.4 of the supplement.

### 2.4. Prior distribution: an improved probabilistic atlas of the thalamus

In Iglesias et al. (2018), we presented a highly detailed probabilistic atlas of the human thalamus built from a combination of *in vivo* MRI and histology. The spatial distribution of the thalamic nuclei was learnt from manual delineations drawn on 3D reconstructed histological sections from 12 specimens (Fig. 3a), whereas 39 MRI scans with manual delineations (Fischl et al., 2002) were used to learn the distribution of surrounding tissue (Fig. 3b). Direct use of this atlas (Fig. 3d) in our new framework is not ideal, as the cerebral white matter was modelled using only two classes – one per hemisphere. While such a parsimonious model with a single component is adequate for modelling the unimodal distribution of white mater intensities in sMRI, it is largely insufficient to model the dMRI orientations. The distribution over white matter voxels is highly multimodal due to the variety of fibre tracts that traverse this tissue in different orientations.

**Figure 3:**
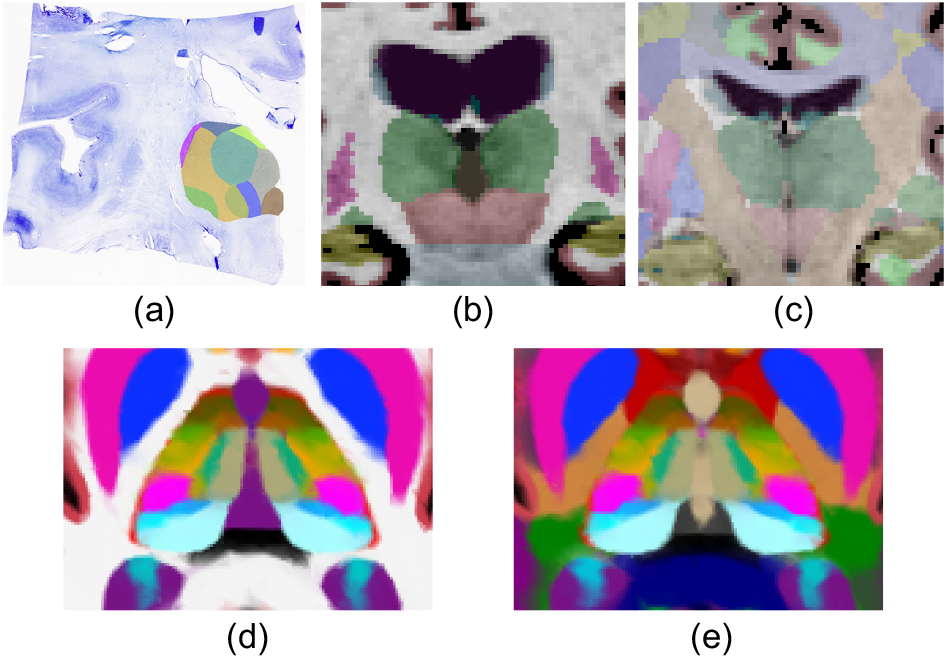
(a-c) Types of segmentations used to build the atlas. (a) Coronal histological section of the thalamus, with manual delineations of the nuclei. (b) Coronal slice of an *in vivo* T1-weighted MRI scan, with manual delineations for whole brain structures. (c) Similar coronal slice of one of the new 16 cases, with the white matter subdivided into tracts. (d-e) Corresponding axial slices of the previous and updated probabilistic atlases. The original atlas (d) was trained with segmentations like the ones in (a-b), while the new atlas used (a-c).

In principle we could model such a complex distribution using a mixture model with many components. However, such an approach is likely to fail, as some of these components may end up modelling non-white-matter tissue. Instead, we have refined our atlas by subdividing the white matter surrounding the thalamus into 45 tracts. To achieve this, we complemented the training data in Iglesias et al. (2018) (12 *ex vivo* thalami and 39 *in vivo* whole brains) with *in vivo* sMRI/dMRI data from 16 additional subjects, that were labelled manually as part of an update (Maffei et al., 2021) to the TRACULA (TRacts Constrained by UnderLying Anatomy) package distributed with FreeSurfer (Yendiki et al., 2011).

The TRACULA training set (16 healthy adults from the publicly available MGH-USC HCP; Fan et al. 2016) consisted of dMRI data, acquired using 512 directions at a maximum b-value of 10,000 s/mm^2^ with 1.5 mm isotropic spatial resolution, and sMRI T1-weighted data, acquired with an MPRAGE sequence at 1 mm isotropic resolution. Cortical parcellations and subcortical segmentations, including the whole thalami and cerebral white matter (left and right), were obtained through FreeSurfer (Dale et al., 1999; Fischl et al., 1999, 2002, 2004). Whole-brain probabilistic tractograms were generated for each subject using constrained spherical deconvolution approaches (Tax et al., 2014; Jeurissen et al., 2014) and streamlines used to manually label 42 white matter tracts through a combination of inclusion and exclusion criteria (Maffei et al., 2021). Resulting tractograms were transformed to the sMRI of the subject using a boundary-based, affine registration method (Greve and Fischl, 2009) and converted into visitation maps. These soft segmentations were spatially smoothed with a Gaussian kernel (*σ* = 2mm). For each white matter voxel in the FreeSurfer subcortical segmentation, we replaced its label by the tract with the highest probability (unless such probability was below 5%), dividing the white matter into 42 tracts and a generic white matter class (Fig. 3c).

The three types of segmentations (Fig. 3a-c) were used to rebuild the atlas, using a technique that enables combining labellings with different levels of detail (Iglesias et al., 2015). The new atlas (Fig. 3e) is almost identical to the old one (Fig. 3d), but now includes more specific subclasses in the white matter. As a last adjustment, we subdivided the anterior commissure and other tracts comprising the corpus callosum, while excluding tracts not passing adjacent to the thalamus. This resulted in 45 final labels for the white matter tracts. Each of these subclasses can be modelled either with unimodal distributions or mixtures with very few components, effectively preventing the modelling of non-white-matter tissue.

### 2.5. Implementation details

#### 2.5.1. Data preparation

We assume that the sMRI has been processed with FreeSurfer, which yields a bias field corrected image and a whole brain segmentation (*aseg*.*mgz*, Fischl et al. 2002). The labels in aseg.mgz are used to initialise both the atlas deformation (Iglesias et al., 2015, 2018) and hyperparameters in the structural prior in Eq. (14). In practice the hypermean 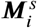 is estimated from the median of the relevant label in this initial coarse segmentation, and 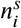 relates to the number of voxels used in estimating 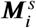. However, it is more difficult to robustly inform prior distributions of the covariance, so we set both 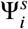 and 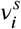 to zero to provide a non-informative prior, giving the set of prior parameters 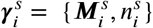.

We also assume that the source dMRI has been put through the preprocessing stages of TRACULA (Yendiki et al., 2011; Maffei et al., 2021). This includes FSL’s eddy current and subject motion correction (Andersson and Sotiropoulos, 2016) before fitting the tensor model. Additionally, we identify DTI voxels with poor fits as those with tensors that have negative eigenvalues or FA outside the range [0, 1]. These are replaced by a local average tensor constructed by convolution of the log space tensors with a 3D Gaussian kernel. These cleaned tensors are converted to the log domain (Arsigny et al., 2006) before interpolation to the voxel grid of the sMRI.

#### 2.5.2. Mixture model specification

The assignment of component distributions to label classes is one of the modelling choices that must be made before segmentation. We assign structural and diffusion components independently for each label class, defining what we will call the structural mixture model (*sMM*) and diffusion mixture model (*dMM*) respectively. In practice, this constrains most weights *g*_*c,i*_ and *w*_*c,j*_ to 0 or 1, with a single component distribution often shared between groups of labels. However, we do allow for many-to-many relationships between the label-classes and components. For example, allowing the structural appearance of the CSF label to be modelled by two Gaussian components, one for “clean” CSF that is also used to model ventricle labels and one for “messy” CSF that is shared with the choroid plexus.

For class likelihoods composed of multiple distributions, the non-zero weights are set to be equal for the first E step and initial component parameters are obtained by use of k-means clustering. Details of this clustering for each likelihood formulation can be found in Section S.3 of the supplement, while optimisation of the default sMM and dMM definitions is performed in Section 3.2.

#### 2.5.3. Reflective symmetry

Another regularising constraint we impose on our segmentation is reflective symmetry of contralateral structures. Classes in one hemisphere share structural distributions with the corresponding classes in the opposite hemisphere. However, in dMRI we assume the average ellipsoids described by tensors from two contralateral structures should be reflections of each other in the median plane. As the head is never positioned in a perfect alignment with the scanner coordinate system, we optimise for a vector normal to the plane of reflection, ***r***, initially assumed to be parallel to the left-right axis of the voxel grid. Prior to the M step, we substitute reflected distribution parameters to the bound in Eq. (8) and formulate the contribution of ***r***, producing an objective function that is fourth order in ***r*** with known first and second derivatives. This objective can be optimised using an interior-point method under the constraint that ‖***r***‖ = 1. We then jointly fit parameters for corresponding component distributions in the left and right hemispheres. Detailed formulations for the reflection optimisation and joint distribution fitting can be found in Section S.1 of the supplement.

#### 2.5.4. Likelihood adjustment

Our model assumes that the resolutions of the dMRI and sMRI are identical. While datasets such as the HCP deviate from this assumption to a lesser degree, conventional quality datasets have much lower resolution for the dMRI in particular, for example T1-weighted images are typically acquired with each voxel dimension at approximately 1 mm while dMRI voxel dimensions can approach 2.5 mm in each direction. As we resample to the resolution of the sMRI, more dMRI voxels are used in likelihood parameter estimation than are available from the source imaging, which leads to overfitting of the dMRI. In practice, we counteract this effect by downplaying the weight of the dMRI voxels in the objective function by a factor *ϵ* equal to the ratio of voxel sizes between dMRI and sMRI. Further details can be found in Section S.2 of the supplement.

## 3. Experiments and Results

To quantitatively evaluate our method and compare between the three likelihood formulations we performed experiments using co-registered sMRI and dMRI from three datasets. In Section 3.1 we generate a population template from HCP subjects, and use it to identify manually segmentable labels corresponding to groups of labels from our histological atlas. In Section 3.2 we use this template to tune our method in a process of model selection. In Section 3.3 we evaluate application of our method to high resolution dMRI on subjects from HCP, including comparisons to manual segmentations and test-retest reliability. Finally, in Section 3.4 we evaluate application of our method to conventional quality dMRI. This includes test-retest reliability on images acquired locally at the University College London Dementia Research Centre (*UCL DRC*) and indirect evaluation on subjects with underlying pathologies by testing our method’s ability to distinguish between healthy controls and subjects with AD from the ADNI dataset.

In the following experiments, when comparing regions of interest (ROIs) corresponding to the same label in two separate segmentations we use the Dice Similarity Coefficient (*DSC*) and 95^th^ percentile of Hausdorff distance (*95HD*). For two ROIs *X* and *Y* these are defined as

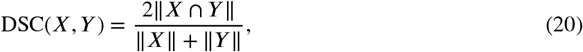

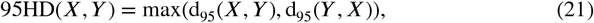

where ‖ ·‖ indicates the volume of the ROI and d_95_(*X, Y*) is the 95^th^ percentile of the set of distances between points on the ROI boundaries, 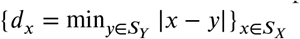.

### 3.1. Population template and manual labels

When evaluating segmentation methods for medical images, it is common practice to compare the resulting label maps to a gold standard, usually obtained from manual delineation by a trained rater. However, manual delineation of 52 histological labels on *in vivo* MRI is infeasible, as many of the boundaries between are invisible at ∼1mm resolution. Manual segmentation protocols for larger groups of thalamic regions (with fewer labels) exist in the literature (Tourdias et al., 2014), but their anatomical definitions are incompatible with those of our histological labels, introducing bias and preventing direct and fair comparison. In this study, our goal is to compare the performance of our tool with a gold standard that is based on our 52 histological labels and informed by both sMRI and dMRI contrast. For this reason, we adapted these labels to define our own manual segmentation criteria for thalamic labels that can be accurately visualised and segmented on a combination of T1-weighted MPRAGE and directionally-encoded colour FA (*DEC-FA*); when labels of smaller thalamic nuclei were not identifiable from the intensity and contrast of the MRIs, these labels were combined and grouped together, so that the boundaries of the original 52 histological atlas labels can be easily matched and compared.

The first step in defining these criteria was to create a high resolution template using 500 subjects from the WashU-UMN HCP dataset (Van Essen et al., 2013) and an unbiased template construction method (Joshi et al., 2004). We used three channels in the registration: T1-weighted intensity, T2-weighted intensity, and FA. In order to include directional information in the template, we used the final set of registrations to align and average the DTI tensors in the log domain. The resolution of the template is equal to the resolution of the HCP sMRI data, i.e., 0.7mm isotropic. Slices from the template are shown in Fig. 4.

**Figure 4:**
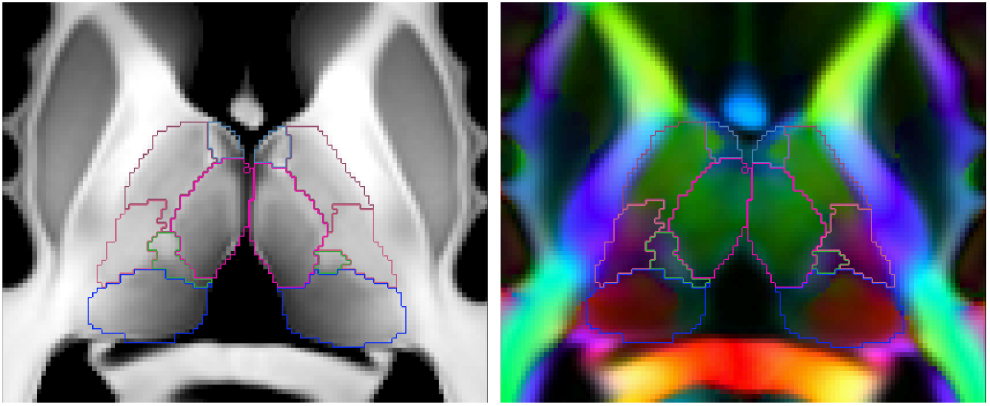
Axial views of the T1-weighted (left) and DEC-FA (right) population templates of the thalamus, overlaid with the outlined labels obtained by manual segmentation. Manually segmented label colour maps are given in Table 1.

As a second step to define the gold standard for comparison, we registered the histological atlas to the template, producing a preliminary segmentation of 52 separate thalamic labels. This preliminary segmentation was then manually refined by an anatomy expert (JA, assisted by MB), to correct any anatomical errors from registration, and to combine those thalamic regions which were not reliably identifiable from the multi-modal template into labels which represent larger thalamic groups. This process resulted in a set of 10 bilateral labels that were manually identifiable from the template. The template labels are used in Section 3.2 to aid in tuning our method, and criteria for these labels are used in Section 3.3 to generate gold standard segmentations for comparison. However, the reduced contrast and resolution of scans for individual subjects resulted in increased ambiguity for some boundaries, hence we further combine our 10 labels into a final set of 5 coarser groupings that are visually identifiable *in vivo*, enabling evaluation without biasing results. Manual labels for the template can be seen in Fig. 4 and the correspondences between the evaluation groupings, manually segmented labels and original histological atlas labels can be seen in Table 1.

**Table 1.**
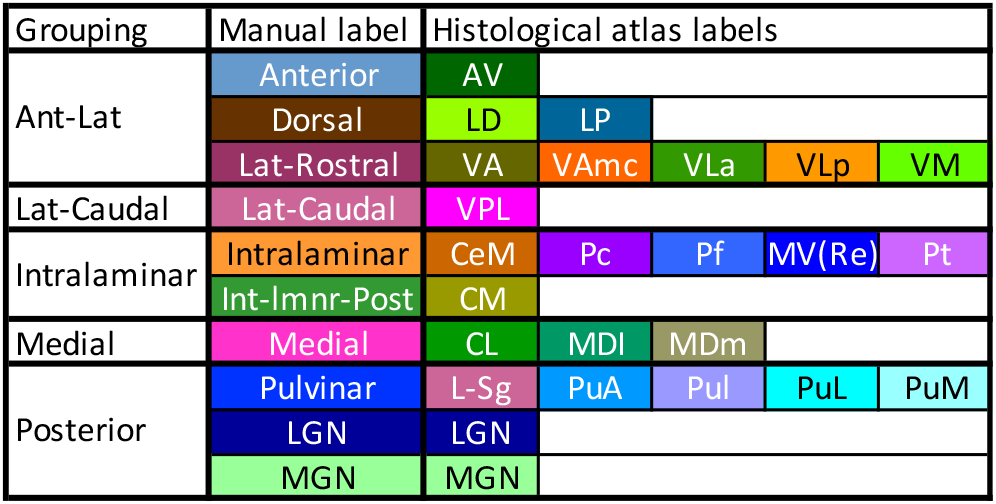
Summary of the label merging operations used to generate the manually segmented labels from histological atlas nuclei, and groupings of manual labels used for evaluation. Displayed colours follow the convention used in figures throughout this manuscript. Abbreviation definitions are listed in Section S.4 of the supplement.

### 3.2. Model selection

Practical implementation of the proposed framework requires decisions on how to share the sMM and dMM parameters (Section 2.5.2), which amounts to a model selection problem. In principle, our generative models enables the computation of the so-called model evidence, which enables comparison of models with different number of parameters. While theoretically appealing, computing this evidence requires marginalisation over all parameters, which leads to intractable integrals that require approximations. Instead, we selected the sMM/dMM groupings with a combination of prior knowledge and a systematic approach called “Technique for Order Preference by Similarity to Ideal Solution” (TOPSIS) Behzadian et al. (2012).

#### Structural groupings

In our previous work, we used two Gaussian components to model the contrast difference between medial and lateral classes (Iglesias et al., 2018). Here, we added a third Gaussian modelling the medial portion of the medial pulvinar (PuM) nucleus, which has a structural appearance closer to grey matter compared with the lateral portion of the PuM. We then compared the atlas prior and histograms of the template volumes to identify 33 possible sMMs grouping nuclei into three component distributions, which were considered by TOPSIS (detailed below).

#### Diffusion groupings

In Section 3.1 we defined 10 labels for each thalamus that are manually identifiable from combined sMRI and DEC-FA. However, inspection of the dMRI tensors within these regions found greater heterogeneity in some regions than in others. As additional borders within these labels could not be confidently matched with boundaries in the histological atlas, we examined multiple options for combining histological nuclei into larger structures to be fit with a component distribution. Including these additional boundaries, and allowing for the possibility of bimodal histograms for some labels, we arrived at 21 possible dMMs, grouping nuclei into between 11 and 13 component distributions.

#### TOPSIS

To optimise the choice of sMMs and dMMs in a systematic fashion, we tested each possible combination of sMM and dMM parameter groupings on the population template. We then evaluated these models by comparing Dice scores and 95HD for the whole thalamus as well as the “grouping” and “manual label” regions listed in Table 1. We then used TOPSIS to create a single, normalised fitness score for each combination of shared parameter specifications allowing them to be ranked. The resulting scores for each likelihood model are shown in Fig. 5. The chosen models are provided in a spreadsheet in the supplementary material as well as descriptions of all candidate models.

**Figure 5:**
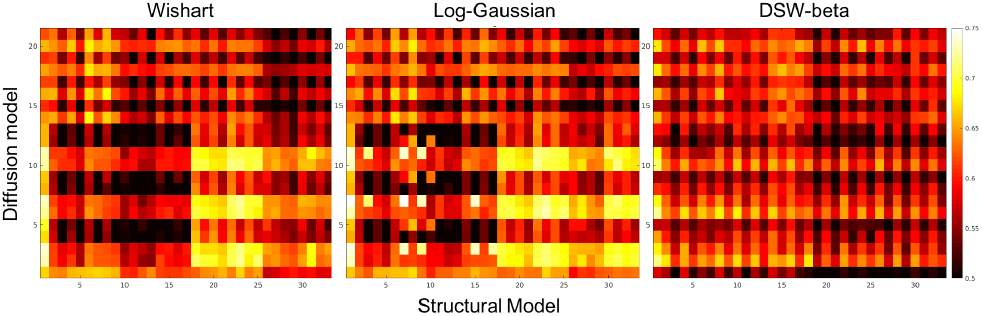
TOPSIS fitness plots for combinations of structural (horizontal axis) and diffusion (vertical axis) grouping models. Plots are displayed for Wishart, Log-Gaussian and DSW-beta likelihood models. A mapping from model numbers to parameter groupings is provided as a spreadsheet in the supplementary material.

### 3.3. Application to high resolution dMRI

Having individually tuned the mixture models and defined a manual protocol corresponding to our histological labels, the obvious next step is to assess the performance of our joint segmentation method on HCP quality data. A comparison of our joint segmentation to both the FreeSurfer whole thalamus segmentation (*aseg*.*mgz*) and our previous structural-only method are shown in Fig. 6. This figure shows each segmentation overlaid on both the T1-weighted sMRI and the DEC-FA for two healthy subjects^2^.

**Figure 6:**
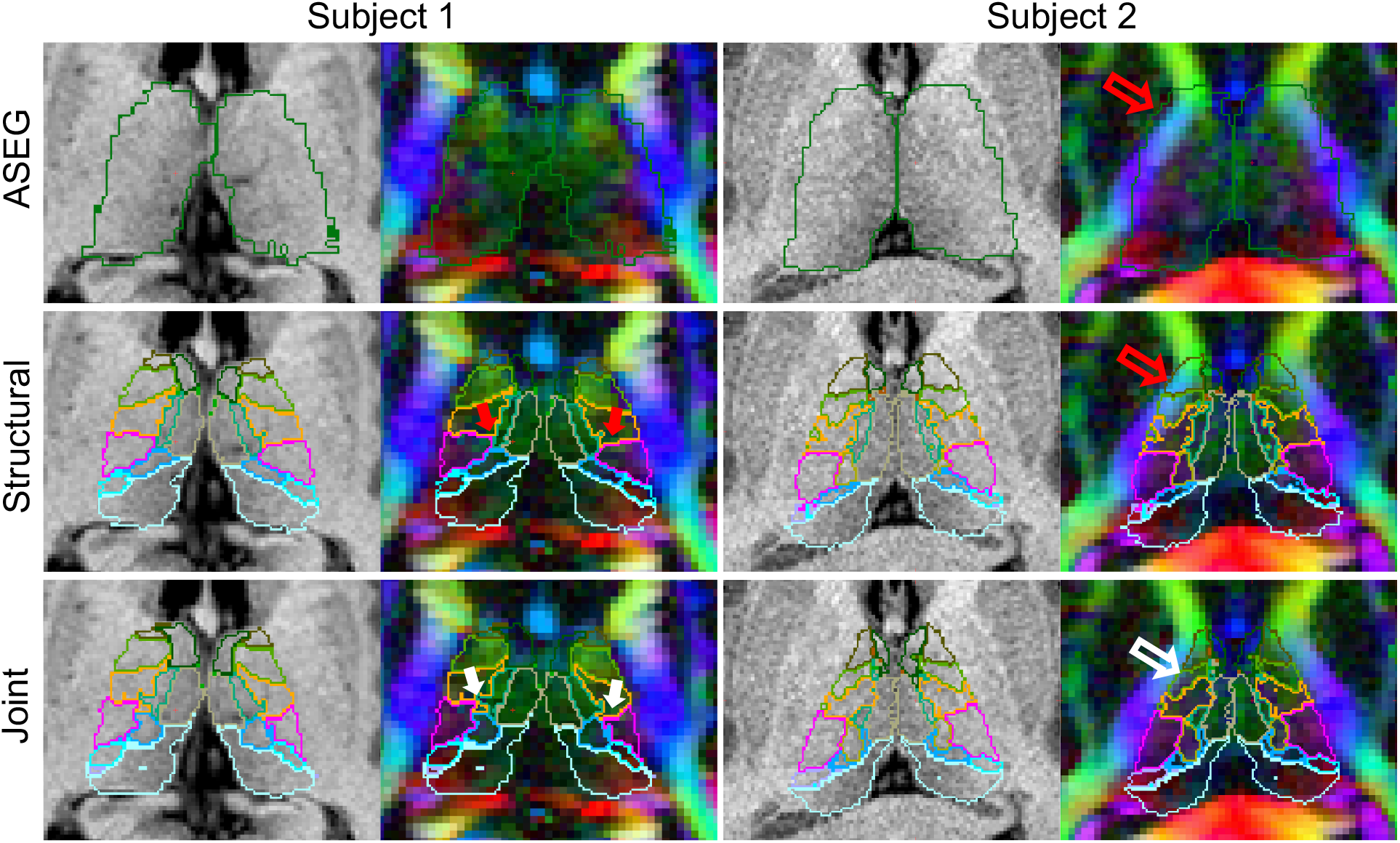
Comparison of thalamic segmentations generated from FreeSurfer’s recon-all (*aseg*.*mgz*), structural and joint (DSW-beta) Bayesian segmentation on two HCP subjects. Arrows have been overlaid to indicate significant instances of correct (white) and incorrect (red) identification of both internal (solid) and external (outline) boundaries.

In both subjects the whole thalamus aseg segmentation, used as an initialisation for both Bayesian methods, shows obvious errors when overlaid on the DEC-FA, with more extreme over-segmentation for subject 2. In subject 1 the structural-only segmentation appears to compensate for these errors and provides an improved exterior boundary. However, our joint method shows marked improvement in the agreement of internal boundaries with colours displayed in the dMRI (solid arrows) as well as a smaller improvement in the exterior boundary. This effect is much more pronounced in subject 2, where the initial over-segmentation of the thalamus propagates to the structural-only method but is corrected by the joint method (arrow outlines).

Such observations provide compelling qualitative evidence for the efficacy of our new method. However, to fully evaluate its usefulness we must quantitatively assess both accuracy and repeatability.

#### 3.3.1. Direct evaluation with manual ground truth

To provide a quantitative measure of segmentation quality, our anatomy expert (JA, assisted by MB) manually segmented images for 10 randomly selected subjects from the WashU-UMN HCP dataset (Van Essen et al., 2013) using the protocol outlined in Section 3.1. The manual segmentations were performed using a combination of T1-weighted and DEC-FA at a 1.25 mm isotropic resolution, corresponding to the native resolution of the diffusion data in HCP. We generated segmentations for these subjects using each of the three joint likelihood implementations from Section 2.3 as well as our previously published structural-only implementation^3^ (Iglesias et al., 2018). These automated segmentations, which have the resolution of the structural scans (0.7 mm), were resampled to 1.25 mm isotropic resolution and compared with the ground truth using DSC and 95HD. Dice scores and 95HD for the five groupings (in column one of Table 1) and the whole thalamus are shown in Fig. 7.

**Figure 7:**
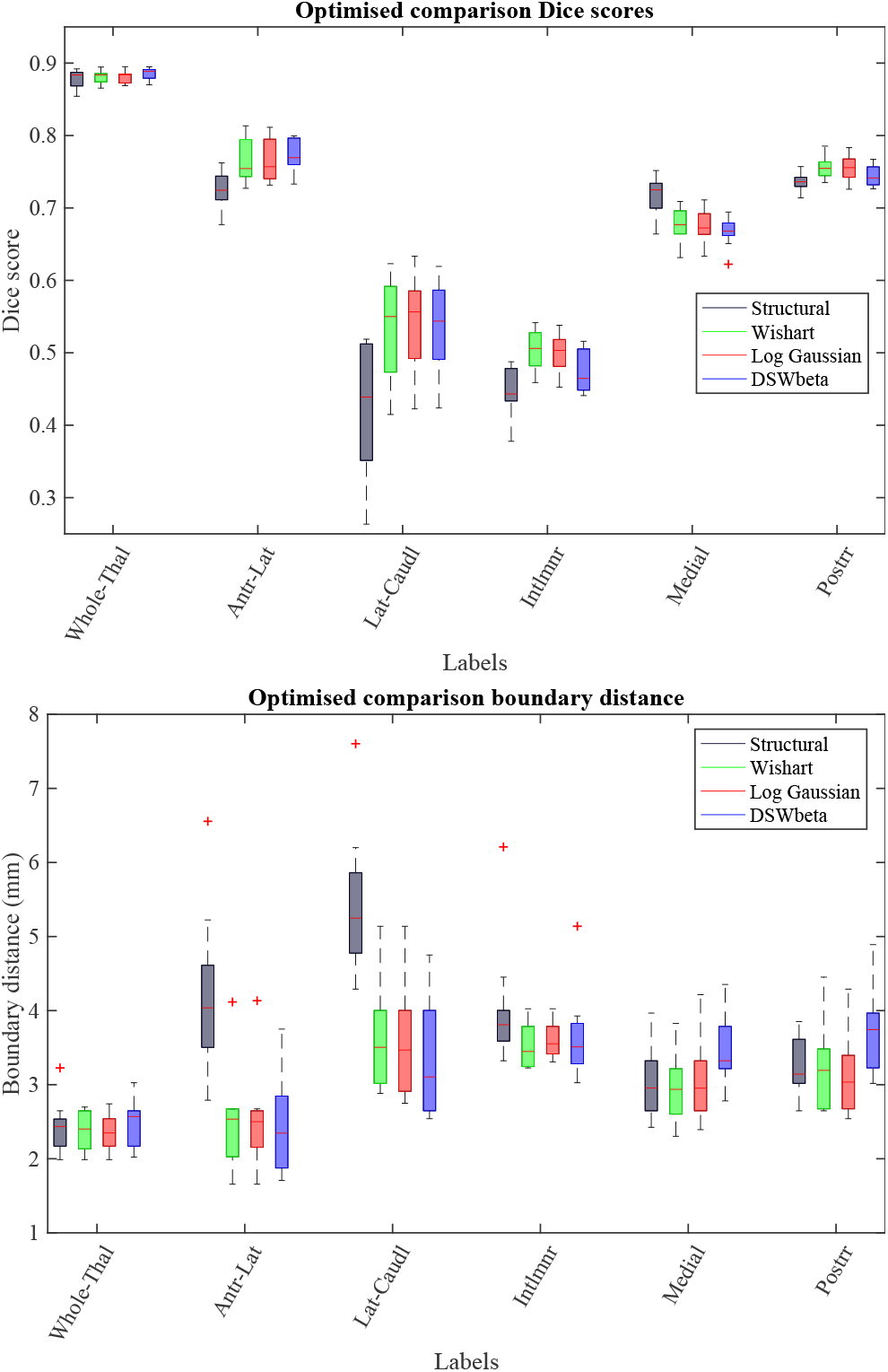
Dice score (top) and 95HD (bottom) comparison of automated thalamic segmentations to manual delineations of 10 HCP subjects. Scores are stated for our previous structural only method as well as the three likelihood implementations of our joint method.

In general, both the DSC and 95HD plots follow similar trends. The median Dice scores for the whole thalamus in structural-only, Wishart and Log-Gaussian implementations were 0.88 with a small increase to 0.89 for DSW-beta. Similarly the 95HD for all methods was between 2.3 and 2.5 mm or equivalent to approximately 2 voxels on the manual segmentations. This contrasts to the marked improvement in the exterior boundary for subject 2 in Fig. 6. As subject 2 was not selected for manual segmentation, the direct comparisons in Fig. 7 indicate that joint segmentation may only provide a small improvement in exterior boundary where the structural segmentation has worked well, but that it is more robust to errors in initialisation and atypical cases.

Of more interest are the interior boundaries. In nearly all labels the joint methods outperform structural-only with lateral-caudal class showing an improvement of 10 Dice points. This can be seen in Fig. 6 where the solid arrows indicate this changes in this boundary for subject 1. The only label class where the structural method outperforms our joint implementation is the medial class. This is expected as a medial-lateral contrast change is modelled explicitly in the structural-only method. However, the difference is small with a median DSC of 0.72 in structural compared to 0.67 for the joint methods and comparable 95HD measurements in this class.

There is comparatively little difference between the three diffusion likelihood implementations. The Wishart and Log-Gaussian implementations show the most similar results, while in the DSW-beta implementation small decreases in accuracy of the intralaminar and posterior classes are offset by improvements in the antero-lateral classes and whole thalamus exterior.

#### 3.3.2. Test-retest reliability analysis

In order to assess the test-retest reliability of the method (a crucial feature in large scale, multi-centre studies), we segmented images from 110 HCP subjects using two different sets of DTI images for each subject – one based on the b=1000 s/mm^2^ shell and one based on the b=2000 s/mm^2^ shell – and compared the outputs. While the results of such an experiment are optimistic when compared to experiments in which images are acquired with multiple scanners, it does enable thorough comparison within the same dataset; test-retest experiments with multiple acquisitions are described in Section 3.4.1 below.

Dice comparison of each likelihood implementation on these two sets of reconstructed tensors can be seen in Fig. 8. These results generally show that all three models are reasonably robust to such an acquisition change in HCP quality data, with a median Dice score of 0.85 or greater in each grouped label across all models. Similarly, each model showed Dice scores of greater than 0.95 for whole thalamus outlines. While median Dice scores for each grouped label are comparable between models, the DSW-beta does appear to be more robust for whole thalamus as well as three of the five grouped labels. Additionally, both Wishart and Log-Gaussian show a larger number of low Dice outliers with minimum values reaching as low as 30 points bellow the median in some cases.

**Figure 8:**
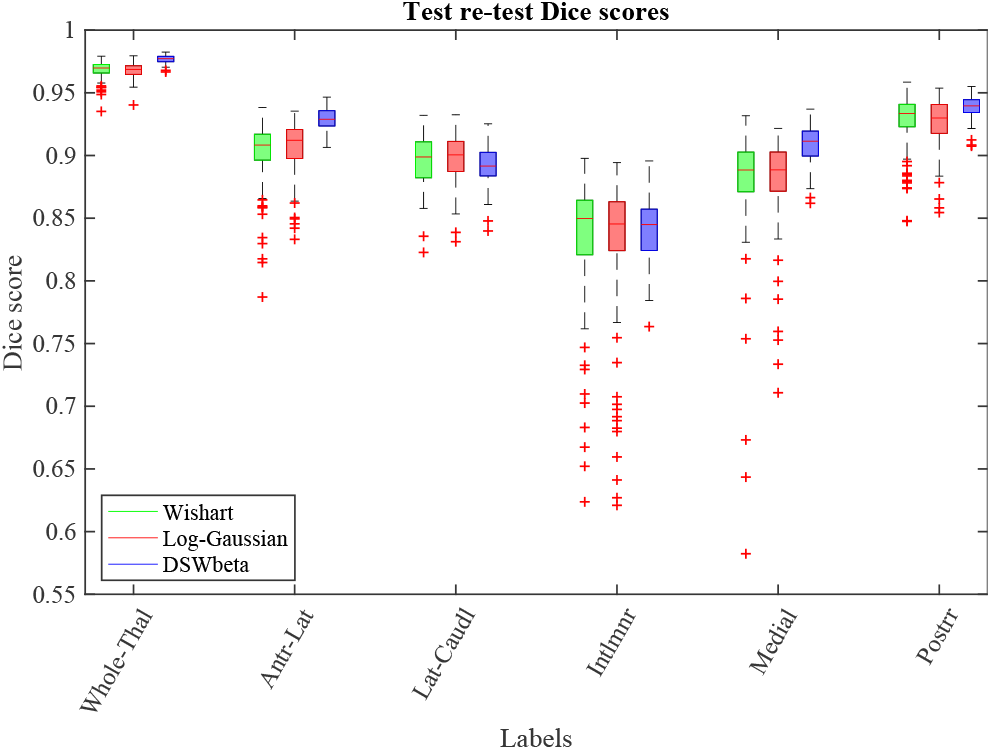
Dice score evaluation of test-retest reliability on 110 HCP subjects. For each subject, we performed two segmentations using DTI images obtained by fitting the tensor to the data from b=1000 s/mm^2^ and b=2000 s/mm^2^ shells separately and computed Dice scores for groups of labels in the two resulting segmentations.

### 3.4. Applications to conventional quality dMRI

While our method assumes that the resolution of the diffusion MRI approaches 1 mm isotropic (which is the case for many modern datasets, e.g., following the HCP protocol), it is of high interest to segment the thalamic nuclei in lower resolution scans for two reasons. First, because large amounts of legacy data were acquired at lower resolution. And second, because many current studies (e.g., ADNI, GENFI) still use those acquisitions, either in order not to deviate from the protocol used to acquire images earlier in the project or to accommodate acquisition constraints such as available scanner time. As explained in Section 2.3 above, compatibility with conventional quality data is actually the reason why we chose to model the diffusion tensor in our likelihood term, rather than using a more sophisticated, higher order model.

Reduced resolution and contrast on such scans, compared to HCP, make manual delineation using the criteria from Section 3.1 infeasible. For this reason we do not do a direct comparison of our methods to the 10 labels defined by manual segmentation. Instead, we first evaluate the reliability of the joint segmentation method through test-retest analysis, then assess the utility of our method, using the ability to discriminate individuals with AD vs controls as a proxy for accuracy.

#### 3.4.1. Test-retest reliability analysis

In order to assess the test-retest reliability of the method on lower resolution dMRI, we used a separate dataset, comprising 21 healthy volunteers (9 male, 12 female, aged 53 – 80 years) acquired at the UCL DRC. Three MRI sequences were performed for each subject in a single session: one T1-weighted MPRAGE 1.1 mm isotropic resolution; and two diffusion weighted acquisitions each consisting of 64 gradient directions at a b-value of 1,000 s/mm^2^ and a 2.5 mm isotropic resolution. Using the two dMRI acquisitions as separate tests, segmentations were performed at a 1 mm isotropic resolution in the native orientation of the individual dMRI volumes before being resampled to the native space of the structural volume for calculation of test-retest Dice scores. The Dice scores for this experiment are shown in Fig. 9.

**Figure 9:**
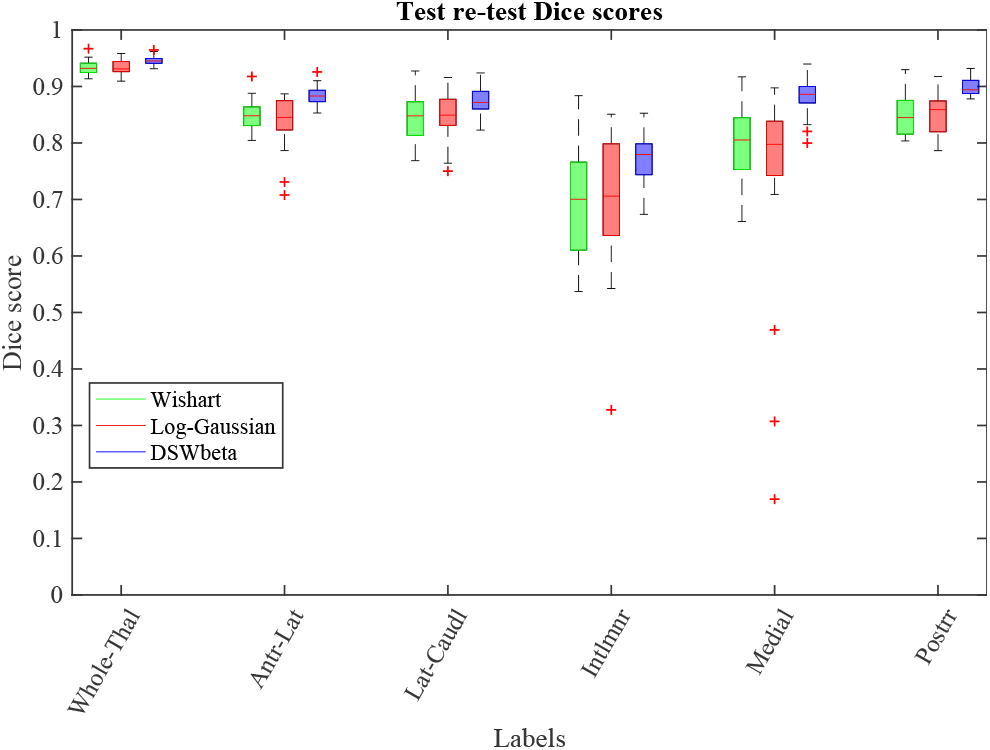
Dice score evaluation of test-retest reliability on conventional-quality data from 21 subjects acquired at the UCL Dementia Research Centre. For each subject, we performed two segmentations using dMRI data acquired in the same session using the same acquisition parameters and computed Dice scores for groups of labels.

While this experiment shows lower Dice scores than in HCP, possibly due to the increased voxel size and reduced quality of the data, median scores are still above 0.9 for whole thalamus and 0.8 for four of the five grouped labels. However, it is more apparent from this plot that both the Wishart and Log-Guassian implementations are less robust in lower quality data. This may be due the increased dimensionality of these two models, meaning imprecise fitting of the tensor model caused by partial volume effects has a greater impact than for the more robust FA and principle direction model used by the DSW-beta likelihood.

#### 3.4.2. Alzheimer’s disease study

In order to evaluate the usefulness of our method in a classical group study with imaging data of conventional quality, we ran a further experiment using the ADNI dataset. The ADNI was launched in 2003 by the National Institute on Aging, the National Institute of Biomedical Imaging and Bioengineering, the Food and Drug Administration, private pharmaceutical companies and non-profit organizations, as a $60 million, 5-year public-private partnership. The main goal of ADNI is to test whether MRI, positron emission tomography (PET), other biological markers, and clinical and neuropsychological assessment can be combined to analyze the progression of MCI and early AD. Markers of early AD progression can aid researchers and clinicians to develop new treatments and monitor their effectiveness, as well as decrease the time and cost of clinical trials. The Principal Investigator of this initiative is Michael W. Weiner, MD, VA Medical Center and University of California — San Francisco. ADNI is a joint effort by co-investigators from industry and academia. Subjects have been recruited from over 50 sites across the U.S. and Canada. The initial goal of ADNI was to recruit 800 subjects but ADNI has been followed by ADNI-GO and ADNI-2. These three protocols have recruited over 1500 adults (ages 55–90) to participate in the study, consisting of cognitively normal older individuals, people with early or late MCI, and people with early AD. The follow up duration of each group is specified in the corresponding protocols for ADNI-1, ADNI-2 and ADNI-GO. Subjects originally recruited for ADNI-1 and ADNI-GO had the option to be followed in ADNI-2. For up-to-date information, see http://www.adni-info.org.

Specifically, we repeated the experiment from our previous article (Iglesias et al., 2019) using the T1-weighted and dMRI scans of 92 subjects from ADNI2. Here we considered 47 subjects with AD and 45 age-matched controls (73.8±7.7 years; 44 females total). The data consisted of T1-weighted scans, with a resolution of 1.2×1×1 mm (sagittal), and dMRI with a resolution of 1.35×1.35×2.7 mm (axial). We fit the DTI model to the b=1000 s/mm^2^ shell (41 directions), combined with 5 volumes at b=0. Given the test-retest results above and considering the similar levels of accuracy seen between dMRI likelihood models in the ground truth comparisons, we focus on our DSW-beta likelihood model in this experiment.

Initial tests on subjects from the ADNI dataset revealed some cases where the inclusion of the dMRI shifts boundaries in the segmentation due to the lower resolution of the dMRI data (and thus increased partial volume effects). An example is the over-segmentation of the thalamus into the CSF in Fig. 10a. We addressed this by allowing the contribution of the dMRI likelihood term to be reduced in proportion to the ratio between voxel volumes in the sMRI and dMRI volumes (Fig. 10b) as outlined in Section 2.5 and Section S.2 of the supplement.

**Figure 10:**
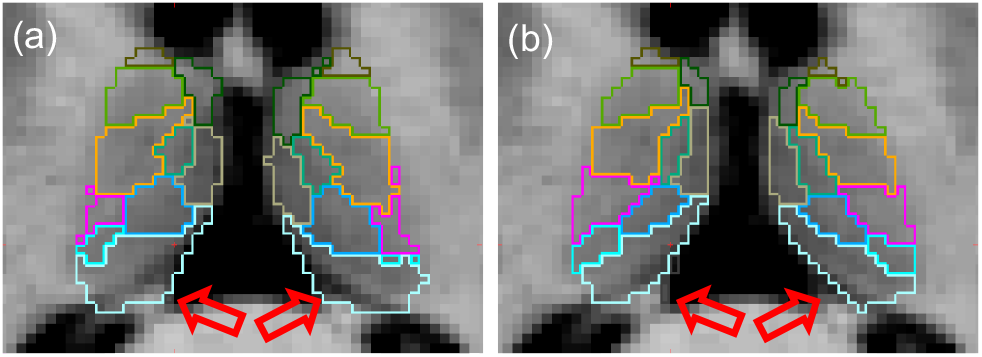
Comparison of thalamic segmentations of a subject from the ADNI dataset using equal (a) and reduced (b) dMRI likelihood weighting. Weighting the dMRI likelihood by the ratio of voxel volumes between sMRI and dMRI results in more accurate estimation of boundaries with heavy partial voluming in the diffusion channel, e.g., the CSF/posterior-thalamus boundary (red arrows).

As in Iglesias et al. 2019, we computed receiver operating characteristic (*ROC*) curves for discrimination of subjects into the two classes using five approaches: three based on thresholding the volume of the whole thalamus (as given by the FreeSurfer recon-all stream, the structural segmentation, and the joint segmentation); and two based on thresholding the likelihood ratio given by a linear discriminant analysis (LDA, Fisher 1936) on the volumes of the histological nuclei (as given by the structural and joint segmentation). The resulting ROC curve is shown in Fig. 11 with the area under the curve (*AUC*), accuracy at the elbow and p-values for comparison of AUC values shown in Table 2.

**Figure 11:**
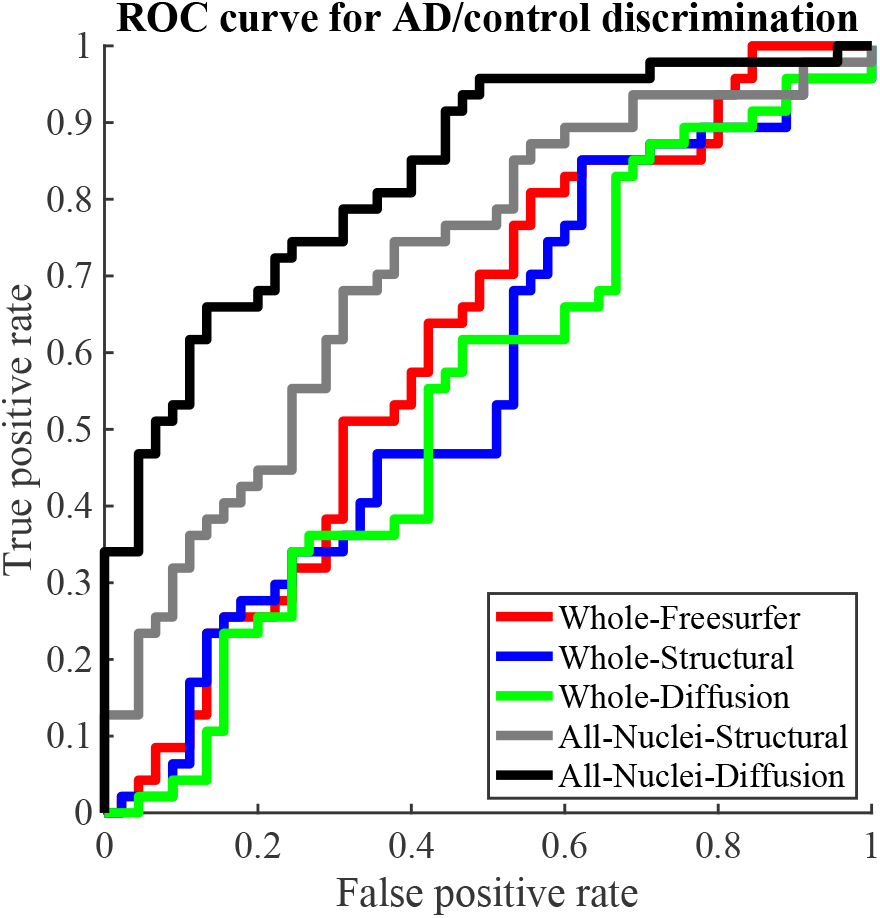
ROC curves for subjects with AD vs controls classification based on thalamic volumes.

**Table 2.** AUC, accuracy at elbow, and p-value for improved AUC values as given by a DeLong test.

|  | FreeSurfer<br>(whole) | Structural<br>(nuclei) | Diffusion<br>(nuclei) |
| --- | --- | --- | --- |
| AUC | 61.28% | 71.30% | **83.36%** |
| Acc. at elbow | 61.96% | 68.48% | **76.09%** |
| p-value vs FreeSurfer |  | 1.6e-01 | **1.1e-03** |
| p-value vs Structural |  |  | **2.1e-02** |

From these curves we can see that all three methods relying on the total volume of the thalamus have poor discriminative ability, with little difference between using FreeSurfer, structural or joint segmentations. This contrasts to the nuclei specific methods, which both show marked improvements. Structural showing an increase of 10% AUC over FreeSufer’s whole thalamus and joint segmentation an increase of 22%. However, only the improvements of the joint method show statistical significance with *p* = 1.1*e* −03 vs. Freesurfer and *p* = 2.1*e* − 02 vs structural nuclei segmentation.

## 4. Discussion and Conclusion

In this article, we have presented and tested a novel segmentation method for thalamic subregions from structural and diffusion MRI. Building on the Bayesian segmentation literature, we use novel likelihood models to exploit structural and diffusion MRI information *jointly* in order to obtain an accurate parcellation of the thalamic nuclei. The information in structural MRI enables placement of boundaries in regions with strong contrast (e.g., medial boundary with the ventricles) with high precision, attributed to its higher resolution; the diffusion information enables the accurate segmentation of boundaries that are invisible in typical structural MRI sequences. Furthermore, we have presented an improved version of our previous histological atlas, which enables more accurate modelling of diffusion MRI in the cerebral white matter. The proposed method will be distributed with FreeSurfer and is widely applicable because the likelihood: *(i)* relies on a simple DTI model, which makes it compatible with virtually every diffusion dataset; *(ii)* adjusts to different resolutions by correcting for voxel sizes; and *(iii)* relies on an unsupervised model that is robust against changes in MR contrast.

We have conducted extensive experiments with manual segmentations, test-retest acquisition, and group studies – including datasets with different resolutions. The results have shown that the joint model exploiting the diffusion information improves accuracy over structural-only segmentation. Moreover, we have also found that the varying resolution gap between structural and diffusion MRI may be accommodated by weighting the diffusion likelihood term to account for voxel size differences, thus bypassing the need to explicitly model partial voluming – which quickly becomes intractable, particularly in multi-modal images defined on different voxel grids. While both our proposed likelihood model (DSW-beta) and the two competing alternatives showed similar levels of improved accuracy over structural-only segmentation, we found the DSW-beta distribution to have the highest test-retest reliability and to be the most robust to changes in the DTI resolution. Compared with other approaches, we produce Dice scores that are within an expected range. For example Su et al. (2019) reported mean scores of 0.64 and above, but direct comparison is complicated by differences in label definition, acquisition type and image resolution.

Our proposed method has a large number of design choices, particularly linked to the specification of shared parameters across classes in the structural and diffusion mixture models. We set these parameters with the combination of expert prior knowledge, a labelled template, and a well-known approach from the decision making literature (TOPSIS). While this approach is suboptimal (our prior knowledge is imperfect; a single template is biased towards a certain population, contrast, and resolution; and TOPSIS’s criteria may not necessarily be ideal), it yielded groupings that worked well in practice for different datasets with different resolution.

This work has two main limitations. First, the lack of quantitative validation of our adapted manual segmentation against other segmentation criteria for the thalamus, e.g., with intra- and inter-rater variability. And second, the lack of explicit modelling for the partial volume effect; while accounting for the voxel size ratio mitigated this problem in our experiments, it is possible that it does not suffice for more extreme ratios. Addressing these two issues with further experiments and solutions based on CNNs remain as future work.

The presented method will be publicly available in FreeSurfer as an extension of our current structural-only code. As high-resolution diffusion data become increasingly accessible, algorithms that can exploit them to produce accurate segmentations – particularly for boundaries that are invisible in structural MRI – have the potential to greatly enhance neuroimaging studies.

## Supporting information

Model selection spreadsheet

## Acknowledgments

This work was primarily funded by Alzheimer’s Research UK (ARUK-IRG2019A003). PG’s work in this area was supported by NIH NIBIB NAC P41EB015902 AY’s work in this area was supported by NIH grants R01 EB021265 and R56 MH121426. DCA’s work in this area was supported by EPSRC grant EP/R006032/1 and Wellcome Trust award 221915/Z/20/Z. The Dementia Research Centre is supported by Alzheimer’s Research UK, Alzheimer’s Society, Brain Research UK, and The Wolfson Foundation. This work was supported by the National Institute for Health Research (NIHR) Queen Square Dementia Biomedical Research Unit and the University College London Hospitals Biomedical Research Centre, the Leonard Wolfson Experimental Neurology Centre (LWENC) Clinical Research Facility, and the UK Dementia Research Institute, which receives its funding from UK DRI Ltd, funded by the UK Medical Research Council, Alzheimer’s Society and Alzheimer’s Research UK. MB is supported by a Fellowship award from the Alzheimer’s Society, UK (AS-JF-19a-004-517). MB’s work was also supported by the UK Dementia Research Institute which receives its funding from DRI Ltd, funded by the UK Medical Research Council, Alzheimer’s Society and Alzheimer’s Research UK. JDR is supported by the Miriam Marks Brain Research UK Senior Fellowship and has received funding from an MRC Clinician Scientist Fellowship (MR/M008525/1) and the NIHR Rare Disease Translational Research Collaboration (BRC149/NS/MH). JEI is supported by the European Research Council (Starting Grant 677697, project BUNGEE-TOOLS) and the NIH (1RF1MH123195-01 and 1R01AG070988-01).

The collection and sharing of the ADNI data was funded by the Alzheimer’s Disease Neuroimaging Initiative (National Institutes of Health Grant U01 AG024904) and Department of Defence (W81XWH-12-2-0012). ADNI is funded by the National Institute on Aging, the National Institute of Biomedical Imaging and Bioengineering, and the following: Alzheimer’s Association; Alzheimer’s Drug Discovery Foundation; BioClinica, Inc.; Biogen Idec Inc.; Bristol-Myers Squibb Company; Eisai Inc.; Elan Pharmaceuticals, Inc.; Eli Lilly and Company; F. Hoffmann-La Roche Ltd and affiliated company Genentech, Inc.; GE Healthcare; Innogenetics, N.V.; IXICO Ltd.; Janssen Alzheimer Immunotherapy Research & Development, LLC.; Johnson & Johnson Pharmaceutical Research & Development LLC.; Medpace, Inc.; Merck & Co., Inc.; Meso Scale Diagnostics, LLC.; NeuroRx Research; Novartis Pharmaceuticals Corporation; Pfizer Inc.; Piramal Imaging; Servier; Synarc Inc.; and Takeda Pharmaceutical Company. The Canadian Institutes of Health Research is providing funds for ADNI clinical sites in Canada. Private sector contributions are facilitated by the Foundation for the National Institutes of Health. The grantee is the Northern California Institute for Research and Education, and the study is coordinated by the Alzheimer’s Disease Cooperative Study at the University of California, San Diego. ADNI is disseminated by the Laboratory for Neuro Imaging at the University of Southern California.

## Footnotes

2 The joint segmentation shown here uses our DSW-beta likelihood model and the structural method has been optimised for the HCP dataset by tuning of the stiffness parameter. For a visual comparison of all likelihood models and the default structural segmentation please see the Section S.5 of the supplement.

3 It should be noted that, to ensure a fair comparison of joint and single channel approaches, the mesh stiffness parameter of the structural implementation was modified to match the joint model that had been developed on the HCP dataset. This improved both the DSC and 95HD structural results compared to the default FreeSurfer distribution.

