## Supplementary material for "Accurate Bayesian segmentation of thalamic nuclei using diffusion MRI and an improved histological atlas"

### S.1. Likelihood and prior model equations

In this section we expand formulae used when implementing the likelihood models for both structural and diffusion MRI. These include formulations of the probability density functions, modifications to account for reflective symmetry as well as the formulae and objective functions required to obtain optimal parameters.

#### *S.1.1. Structural likelihood - Gaussian*

The likelihood model for the structural MRI is a mixture of Gaussian distribution components. To make our segmentation more robust we place a Normal-Inverse-Wishart (*NIW*) prior on the component means and variances. This prior distribution was chosen as it is the conjugate prior for a Gaussian.

*Likelihood.* The parameter set characterising the structural component distributions are $\boldsymbol{\theta}_i^s = \{\boldsymbol{\mu}_i, \Sigma_i\}$, where $i$ indexes the structural components. Under this formulation the likelihood model is

$$\begin{aligned} \boldsymbol{s}_v &\sim \mathcal{N}(\boldsymbol{\mu}_i, \Sigma_i), \\ p(\boldsymbol{s}_v|\boldsymbol{\theta}_i^s) &= \frac{\exp\left\{-\frac{1}{2}(\boldsymbol{s}_v - \boldsymbol{\mu}_i)^\top \Sigma^{-1}(\boldsymbol{s}_v - \boldsymbol{\mu}_i)\right\}}{(2\pi)^{k/2}|\Sigma_i|^{1/2}}, \end{aligned} \tag{S.1}$$

where $k$ is number of structural channels (i.e., the number of input structural images) such that $\boldsymbol{s}_v, \boldsymbol{\mu}_i \in \mathbb{R}^k$ and $\Sigma_i \in \mathbb{R}^{k\times k}$.

In the main paper experiments we have $k = 1$ as we primarily use T1-weighted MPRAGE images as our structural channels, although we have explored including T2-weighted imaging as a second channel and found limited benefit. Additionally, we initialise our model with a small number of structural Generalised Expectation Maximisation (*GEM*) iterations in which a low dimensional diffusion image is used to supplement the T1, using a structural likelihood with $k = 2$. For both the Wishart and log-Gaussian methods this low dimensional diffusion image is the log-determinant of the voxel-wise tensors. For the Dimroth-Scheidegger-Watson-beta (*DSW-beta*) method the low dimensional diffusion image used is the fractional anisotropy.

*Corresponding author

*Prior.* The parameter set characterising the prior on structural component $i$ is $\boldsymbol{\gamma}_i^s = \{\boldsymbol{M}_i^s, n_i^s, \Psi_i^s, \nu_i^s\}$. Hence the structural prior distribution is

$$\boldsymbol{\mu}_i, \Sigma_i | \boldsymbol{\gamma}_i^s \sim \mathcal{NIW}(\boldsymbol{M}_i^s, n_i^s, \Psi_i^s, \nu_i^s)$$

$$p(\boldsymbol{\mu}_i, \Sigma_i | \boldsymbol{\gamma}_i^s) = \frac{\exp\left\{-\frac{n_i^s}{2}(\boldsymbol{\mu}_i - \boldsymbol{M}_i^s)^\top \Sigma_i^{-1}(\boldsymbol{\mu}_i - \boldsymbol{M}_i^s)\right\}}{(2\pi)^{k/2}\left|\frac{1}{n_i^s}\Sigma_i\right|^{1/2}} \times \frac{|\Psi_i^s|^{\nu_i^s/2}\exp\left\{-\frac{1}{2}\operatorname{tr}(\Psi\Sigma_i^{-1})\right\}}{2^{k\nu_i^s/2}\Gamma_k\left(\frac{\nu_i^s}{2}\right)|\Sigma_i|^{(\nu_i^s+k+1)/2}}. \tag{S.2}$$

*Parameter estimation.* Following Eq. (10) from the main paper, optimisation of parameters for each structural component distribution $i$ may be performed independently. Substituting Eqs. (S.1) and (S.2) into Eq. (10) and setting derivatives to zero yields closed form solutions for the optimal mean and covariance

$$\hat{\boldsymbol{\mu}}_i = \frac{n_i^s \boldsymbol{M}_i^s + \sum_{v=1}^V q_v^i \boldsymbol{s}_v}{n_i^s + \sum_{v=1}^V q_v^i}, \tag{S.3}$$

$$\hat{\Sigma}_i = \frac{\Psi_i^s + n_i^s(\boldsymbol{\mu} - \boldsymbol{M}_i^s)(\boldsymbol{\mu} - \boldsymbol{M}_i^s)^\top + \sum_{v=1}^V q_v^i(\boldsymbol{s}_v - \boldsymbol{\mu})(\boldsymbol{s}_v - \boldsymbol{\mu})^\top}{\nu_i^s + k + \sum_{v=1}^V q_v^i + 2}, \tag{S.4}$$

where $q_v^i$ are the soft-segmentations corresponding to structural component $i$

$$q_v^i \leftarrow \sum_{c=1}^C \sum_{j=1}^W q_v^{c,i,j}. \tag{S.5}$$

To initialise our prior parameters, we set $\boldsymbol{M}_i^s$ to the median value of an initial coarse segmentation provided by Freesurfer's recon-all stream (*aseg.mgz*) and $n_i^s$ as the volume of component $i$ in that segmentation. However, as it is difficult to characterise the variances of these structural components *a priori*, we generally set both $\nu_i^s$ and $\Psi_i^s$ to zero to provide a non-informative prior on the covariance as in (Iglesias et al., 2018).

#### *S.1.2. Diffusion likelihood - Wishart*

In the Wishart distribution model for the diffusion MRI, we define $\boldsymbol{d}_v$ as the inverse of the diffusion tensor.

*Likelihood.* The parameter set characterising the Wishart component distributions are $\boldsymbol{\theta}_j^d = \{n_j^d, V_j^d\}$, where $j$ indexes the Wishart components. Under this formulation the diffusion likelihood model is

$$\boldsymbol{d}_v \sim \mathcal{W}(n_j^d, V_j^d),$$

$$p(\boldsymbol{d}_v | \boldsymbol{\theta}_j^d) = \frac{|\boldsymbol{d}|^{(n-4)/2}\exp\left\{-\frac{1}{2}\operatorname{tr}(V^{-1}\boldsymbol{d})\right\}}{2^{3n/2}\Gamma_3\left(\frac{n}{2}\right)|V|^{n/2}}, \tag{S.6}$$

where $\operatorname{tr}(\cdot)$ indicates the trace operation.

*Prior.* Fitting the Wishart components without a prior can result in distributions which are heavily peaked around their means ($\bar{\boldsymbol{d}} = n_j^d V_j^d$). In the setting of our Bayesian segmentation this can lead to the algorithm overfitting labels and may lead to instability in the segmentation. To counter this we set a non-informative prior on the degrees of freedom $n_j^d$ to increase the breadth of the resulting distribution (Görür and Rasmussen, 2010). The parameter set characterising the prior on structural component $i$ is $\boldsymbol{\gamma}_j^d = \{\alpha_j, \beta_j\}$ set to 0.5 and 1.5 respectively. The resulting prior distribution is therefore

$$\frac{(n_j^d - 2)}{2} \sim \Gamma(\alpha, \beta)$$

$$p\left(\frac{(n_j^d-2)}{2} | \boldsymbol{\gamma}_j^d\right) = \frac{\beta_j^{\alpha_j}}{\Gamma(\alpha_j)}\left(\frac{(n_j^d - 2)}{2}\right)^{\alpha_j - 1} \exp\left\{-\beta_j \frac{(n_j^d - 2)}{2}\right\}. \tag{S.7}$$

*Parameter estimation.* In this model, optimising the contribution of component $j$ to the objective function in Eq. 11 from the main paper may be reformulated as a one dimensional maximisation problem with respect to the degrees of freedom $n_j^d$. To do this we first define the soft segmentation $q_v^j$ of component $j$ and the mean tensor $\bar{\boldsymbol{d}}$ as

$$q_v^j \leftarrow \sum_{c=1}^{C}\sum_{i=1}^{G} q_v^{c,i,j}, \qquad \bar{\boldsymbol{d}} \leftarrow \frac{\sum_{v=1}^{V} q_v \boldsymbol{d}_v}{\sum_{v=1}^{V} q_v}. \tag{S.8}$$

Substituting these, and the formulae for the prior and likelihood, into Eq. 11 we obtain the optimisation problem

$$\hat{n}_j^d = 2 \times \arg\max_{\nu} \left\{ (\alpha - 1)\log(\nu - 1) - \beta\nu + (\nu - 2)\sum_{v=1}^{V} q_v \log|\boldsymbol{d}_v| + \left[3\nu\log\nu - 3\nu - \nu\log|\bar{\boldsymbol{d}}| - \log\Gamma_3(\nu)\right]\sum_{v=1}^{V} q_v \right\}, \tag{S.9}$$

$$\hat{V}_j^d = \frac{\bar{\boldsymbol{d}}}{n_j^d}. \tag{S.10}$$

*Reflection estimation.* To further enhance the diffusion model, we also incorporate reflective symmetry, which provides enhanced robustness. To do this we assume the average ellipsoids described by tensors from two contralateral structures should be reflections of each other in the median plane. As the head is never positioned in a perfect alignment with the scanner coordinate system, we optimise for a vector normal to the plane of reflection, $\hat{\boldsymbol{r}}$. With this reflection vector, the relationship between the model parameters for two reflected components $j$ and $j'$ are given by

$$R = I - 2\hat{\boldsymbol{r}}\hat{\boldsymbol{r}}^\top, \qquad n_j^d = n_{j'}^d, \qquad V_j^d = RV_{j'}^d R. \tag{S.11}$$

We then define the soft segmentations and mean tensor for the component as

$$q_v^j \leftarrow \sum_{c=1}^{C}\sum_{i=1}^{G} q_v^{c,i,j}, \qquad q_v^{j'} \leftarrow \sum_{c=1}^{C}\sum_{i=1}^{G} q_v^{c,i,j'}, \qquad \bar{\boldsymbol{d}} \leftarrow \frac{\sum_{v=1}^{V}\left[q_v^j \boldsymbol{d}_v + q_v^{j'} R\boldsymbol{d}_v R\right]}{\sum_{v=1}^{V}\left[q_v^j + q_v^{j'}\right]}. \tag{S.12}$$

Substituting these into Eq. 11 and combining the contributions of both reflected components to the bound, we obtain a one dimensional optimisation problem for the shared degrees of freedom for both components.

$$n_j^d = 2 \times \arg\max_{\nu} \left\{ (\alpha - 1)\log(\nu - 1) - \beta\nu + (\nu - 2)\sum_{v=1}^{V}\left[q_v^j + q_v^{j'}\right]\log|\boldsymbol{d}_v| + \left[3\nu\log\nu - 3\nu - \nu\log|\bar{\boldsymbol{d}}| - \log\Gamma_3(\nu)\right]\sum_{v=1}^{V}\left[q_v^1 + q_v^2\right] \right\}. \tag{S.13}$$

The shape parameters $V_j^d$ and $V_{j'}^d$ may then be calculated as

$$V_j^d = \frac{\bar{\boldsymbol{d}}}{n_j^d}, \qquad V_{j'}^d = \frac{R\bar{\boldsymbol{d}}R}{n_j^d}. \tag{S.14}$$

After the likelihood parameters have been updated by GEM we are then able to optimise the reflection vector by substituting the updated parameters into Eq. 8 from the main paper and simplifying using the relationships in Eq. (S.12). This results in an optimisation problem for the reflection vector

$$\hat{\boldsymbol{r}} = \arg\max_{r:\|\boldsymbol{r}\|=1} \left\{ -\frac{1}{2}\sum_{j=1}^{W} \operatorname{tr}\left( \left(V_j^d\right)^{-1} (I - 2\boldsymbol{r}\boldsymbol{r}^\top) \sum_{v=1}^{V}\left[q_v^{j'}\boldsymbol{d}_v\right](I - 2\boldsymbol{r}\boldsymbol{r}^\top) \right) \right\}. \tag{S.15}$$

This equation is fourth order in $\boldsymbol{r}$ with first and second derivatives that may be calculated analytically. We optimise for $\hat{\boldsymbol{r}}$ using an interior point method. In practice we initialise $\boldsymbol{r}$ parallel to the left-right axis, which will be close to the true solution, making it unlikely for the optimisation to encounter a local maximum.

#### S.1.3. Diffusion likelihood - log-Gaussian

In the log-Gaussian distribution model for diffusion MRI, we define $\boldsymbol{d}_v$ as a vector containing the six independent variables in the natural log of the diffusion tensors $T_v$,

$$\boldsymbol{d}_v = P\,\mathrm{vec}(\log T_v). \tag{S.16}$$

Here $\log T_v$ is the natural matrix log of $T_v$, which can be calculated from the eigenvector decomposition provided by FSL's dtifit tool. We use the $6 \times 9$ matrix

$$P = \begin{bmatrix} 1 & 0 & 0 & 0 & 0 & 0 & 0 & 0 & 0 \\ 0 & 1/\sqrt{2} & 0 & 1/\sqrt{2} & 0 & 0 & 0 & 0 & 0 \\ 0 & 0 & 1/\sqrt{2} & 0 & 0 & 0 & 1/\sqrt{2} & 0 & 0 \\ 0 & 0 & 0 & 0 & 1 & 0 & 0 & 0 & 0 \\ 0 & 0 & 0 & 0 & 0 & 1/\sqrt{2} & 0 & 1/\sqrt{2} & 0 \\ 0 & 0 & 0 & 0 & 0 & 0 & 0 & 0 & 1 \end{bmatrix} \tag{S.17}$$

to reduce the vectorised tensor to 6 independent elements as we then have that $||\boldsymbol{d}_v||_2 = ||\log T_v||_2$ and

$$\mathrm{vec}(\log T_v) = P^\top \boldsymbol{d}_v.$$

*Likelihood.* The parameter set characterising the log-Gaussian component distributions is $\boldsymbol{\theta}_j^d = \{\boldsymbol{m}_j^d, \sigma_j^d\}$, which is a vector mean and scalar standard deviation respectively with $j$ indexing the components. Under this formulation the diffusion likelihood model is

$$\boldsymbol{d}_v \sim \mathcal{N}(\boldsymbol{m}_j^d, \sigma_j^d),$$
$$p(\boldsymbol{d}_v|\boldsymbol{\theta}_j^d) = \frac{\exp\left(-\frac{(\sigma_j^d)^{-2}}{2}(\boldsymbol{d}_v - \boldsymbol{m}_j^d)^\top(\boldsymbol{d}_v - \boldsymbol{m}_j^d)\right)}{(2\pi)^3(\sigma_j^d)^6}. \tag{S.18}$$

*Parameter estimation.* Substituting Eq. (S.18) into Eq. (11) from the main paper and setting derivatives to zero yields closed form solutions for the optimal mean and standard deviation

$$\boldsymbol{m}_j^d = \frac{\sum_{v=1}^V q_v \boldsymbol{d}_v}{\sum_{v=1}^V q_v}, \tag{S.19}$$
$$(\sigma_j^d)^2 = \frac{\sum_{v=1}^V q_v(\boldsymbol{d}_v - \boldsymbol{m}_j^d)^\top(\boldsymbol{d}_v - \boldsymbol{m}_j^d)}{6\sum_{v=1}^V q_v}, \tag{S.20}$$

where is the $q_v^j$ soft segmentation of component $j$

$$q_v^j = \sum_{c=1}^C \sum_{i=1}^G q_v^{c,i,j}.$$

*Reflection estimation.* As with the Wishart distribution we are able to constrain the log-Gaussian diffusion model by incorporating reflective symmetry. Again, we optimise for a vector normal to the plane of reflection, $\hat{\boldsymbol{r}}$. With this reflection vector, the relationship between the model parameters for two reflected components $j$ and $j'$ are given by

$$R = P\,[(I - 2\hat{\boldsymbol{r}}\hat{\boldsymbol{r}}^\top) \otimes (I - 2\hat{\boldsymbol{r}}\hat{\boldsymbol{r}}^\top)]\,P^\top, \qquad \sigma_j^d = \sigma_{j'}^d, \qquad \boldsymbol{m}_j^d = R\boldsymbol{m}_{j'}^d. \tag{S.21}$$

We then define the soft segmentations for the two components as

$$q_v^j \leftarrow \sum_{c=1}^C \sum_{i=1}^G q_v^{c,i,j}, \qquad q_v^{j'} \leftarrow \sum_{c=1}^C \sum_{i=1}^G q_v^{c,i,j'}, \tag{S.22}$$

and can retrieve our new parameter estimates

$$\boldsymbol{m}_j^d = \frac{\sum_{v=1}^V \left[q_v^j \boldsymbol{d}_v + q_v^{j'} R \boldsymbol{d}_v\right]}{\sum_{v=1}^V \left[q_v^j + q_v^2\right]} \tag{S.23}$$

$$(\sigma_j^d)^2 = \frac{\sum_{v=1}^V \left[q_v^j \|\boldsymbol{d}_v - \boldsymbol{m}_j^d\|_2^2 + q_v^{j'} \|\boldsymbol{d}_v - R\boldsymbol{m}_j^d\|_2^2\right]}{6 \sum_{v=1}^V \left[q_v^j + q_v^{j'}\right]} \tag{S.24}$$

To fit the reflection we can substitute $\boldsymbol{m}_j^d = R\boldsymbol{m}_{j'}^d$ into Eq. (S.18) and examine the contributions of the reflection tensor $R$ to the bound in Eq. 8 from the main paper. This gives a formula for the bound

$$Q_{\bar{R}} = \sum_{J=1}^{w} (\sigma_j^d)^{-2} \sum_{v=1}^{V} q_v^j \boldsymbol{d}_v^\top R \boldsymbol{m}_{j'}^d + Z[\cdot], \tag{S.25}$$

where $Z[\cdot]$ does not depend on the reflection. This bound can be optimised directly, but by substituting

$$\bar{M}_j^d = (\sigma_j^d)^{-2} \sum_{v=1}^{V} q_v^j \log T_v, \qquad \operatorname{vec} M_j^d = \boldsymbol{m}_j^d$$

we can recover a similar optimisation problem to the one in Eq. (S.15) for which we already know the first and second derivatives. Hence we can recover the reflection by optimising

$$\hat{\boldsymbol{r}} = \underset{r:\|\boldsymbol{r}\|=1}{\arg\max} \left\{ \sum_{j=1}^{W} \operatorname{tr}\left(\bar{M}_j^d (I - 2\boldsymbol{r}\boldsymbol{r}^\top) M_{j'}^d (I - 2\boldsymbol{r}\boldsymbol{r}^\top)\right) \right\} \tag{S.26}$$

where dMRI component $j'$ is a reflection of component $j$ and Eq. (S.26) is constrained by $\boldsymbol{r}^\top \boldsymbol{r} = 1$

#### *S.1.4. Diffusion likelihood - DSW-beta*

In the DSW-beta distribution model for diffusion MRI, we define $\boldsymbol{d}_v$ as the combination of each tensor's fractional anisotropy $f_v$ and its principle direction $\boldsymbol{\phi}_v$

$$\boldsymbol{d}_v = \{f_v, \boldsymbol{\phi}_v\}. \tag{S.27}$$

*Likelihood.* The parameter set characterising the log-Gaussian component distributions is $\boldsymbol{\theta}_j^d = \{\alpha_j^d, \beta_j^d, \boldsymbol{\psi}_j^d, \kappa_j^d\}$, where $\alpha$ and $\beta$ define a Beta distribution on the FA, and $\boldsymbol{\psi}_j^d$ and $\kappa_j^d$ are the mean and concentration of the DSW distribution. Under this formulation the likelihood of $\boldsymbol{d}_v$ is defined as

$$p(\boldsymbol{d}_v|\boldsymbol{\theta}_j^d) = p(f_v|\alpha_j^d, \beta_j^d) p(\boldsymbol{\phi}_v|\psi_j^d, \kappa_j^d f_v) \tag{S.28}$$

$$p(f_v|\alpha_j^d, \beta_j^d) = B(\alpha_j^d, \beta_j^d) f_v^{\alpha_j^d - 1} (1 - f_v)^{\beta_j^d - 1} \tag{S.29}$$

$$p(\boldsymbol{\phi}_v|\psi_j^d, \kappa_j^d f_v) = \left[Z(\kappa_j^d f_v)\right]^{-1} \exp\left\{\kappa_j^d f_v \left((\boldsymbol{\psi}_j^d)^\top \boldsymbol{\phi}_v\right)^2\right\} \tag{S.30}$$

where $B(\alpha, \beta)$ is the beta function

$$B(\alpha, \beta) = \frac{\Gamma(\alpha)\Gamma(\beta)}{\Gamma(\alpha + \beta)}$$

and $Z$ is the Kummer function in 3D

$$Z(\kappa_j^d f_v) = \int_0^1 \exp(\kappa_j^d f_v t^2) \mathrm{d}t.$$

*Parameter estimation.* Substitution of $q_v^j$ and the DSW-beta likelihood into Eq. (11) from the main paper yields three optimisation problems. The $\alpha$ and $\beta$ parameters may be obtained through a single 2D optimisation

$$\{\alpha_j^d, \beta_j^d\} = \underset{\alpha,\beta}{\arg\max} \left\{ (\alpha - 1) \sum_{v=1}^{V} [q_v \log f_v] + (\beta - 1) \sum_{v=1}^{v} [q_v \log(1 - f_v)] - \log B(\alpha, \beta) \sum_{v=1}^{v} q_v \right\}, \tag{S.31}$$

which we solve using conjugate gradients. For the mean direction the resulting optimisation is

$$\begin{aligned} \psi_j^d &= \underset{\psi:\|\psi\|=1}{\arg\max} \left\{ \psi^\top \left[ \sum_{v=1}^{V} q_v f_v \phi_v \phi_v^\top \right] \psi \right\} \\ &= \underset{\psi:\|\psi\|=1}{\arg\max} \left\{ \psi^\top \bar{M} \psi \right\} \end{aligned} \tag{S.32}$$

with closed-form solution given by the leading eigenvector of $\bar{M}$. Finally the concentration optimisation is given by

$$\kappa_j^d = \underset{\kappa}{\arg\max} \left\{ \kappa \sum_{v=1}^{V} q_v f_v (\psi_j^{d\top} \phi_v)^2 - \sum_{v=1}^{V} q_v \log Z(f_v \kappa) \right\} \tag{S.33}$$

which we solve with the conjugate gradient method.

*Reflection estimation.* As with the Wishart and log-Gaussian distributions we are able to constrain the DSW-beta diffusion model by incorporating reflective symmetry. Under this formulation we again optimise for a vector normal to the plane of reflection, $\hat{\boldsymbol{r}}$, which is used to establish a reflection on a subset of the model parameters. In this case we require that for two reflected components $j$ and $j'$ the beta distribution be the same and the concentration of the DSW distributions are equal;

$$\alpha_j^d = \alpha_{j'}^d, \qquad \beta_j^d = \beta_{j'}^d, \qquad \kappa_j^d = \kappa_{j'}^d.$$

We then place the reflection constraint on the mean vector of the DSW distribution for the pair of reflected components with

$$R = I - 2\hat{\boldsymbol{r}}\hat{\boldsymbol{r}}^\top, \qquad \psi_j^d = R\psi_{j'}^d.$$

Under these constraints we follow a similar process of fitting the reflected components and the reflection itself. Again the soft segmentations are defined as

$$q_v^j \leftarrow \sum_{c=1}^{C} \sum_{i=1}^{G} q_v^{c,i,j}. \qquad q_v^{j'} \leftarrow \sum_{c=1}^{C} \sum_{i=1}^{G} q_v^{c,i,j'}.$$

The optimisation for the Beta distribution parameters then becomes

$$\begin{aligned} \{\alpha_j^d, \beta_j^d\} = \underset{\alpha,\beta}{\arg\max} \Bigg\{ &(\alpha - 1) \sum_{v=1}^{V} [(q_v^j + q_v^{j'}) \log f_v] + (\beta - 1) \sum_{v=1}^{v} [(q_v^j + q_v^{j'}) \log(1 - f_v)] \\ &- \log B(\alpha, \beta) \sum_{v=1}^{v} (q_v^j + q_v^{j'}) \Bigg\}. \end{aligned} \tag{S.34}$$

To optimise for the DSW parameters, we then define mean matrices for each class

$$\bar{M}_j = \sum_{v=1}^{V} q_v^j f_v \phi_v \phi_v^\top \qquad \bar{M}_{j'} = \sum_{v=1}^{V} q_v^{j'} f_v \phi_v \phi_v^\top \tag{S.35}$$

leading to formulations of the mean direction

$$\psi_j^d = \underset{\psi:\|\psi\|=1}{\arg\max} \left\{ \psi^\top [\bar{M}_j + R\bar{M}_{j'} R] \psi \right\} \tag{S.36}$$

and concentration

$$\kappa_j^d = \arg\max_{\kappa} \left\{ \kappa \sum_{v=1}^{V} \left[ q_v^j f_v (\boldsymbol{\psi}_j^{d\top} \boldsymbol{\phi}_v)^2 + q_v^{j'} f_v (\boldsymbol{\psi}_{j'}^{d\top} \boldsymbol{\phi}_v)^2 \right] - \sum_{v=1}^{V} (q_v^j + q_v^{j'}) \log Z(f_v \kappa) \right\}. \tag{S.37}$$

Finally, as with the other two tensor likelihood models, we are able to refine the reflection vector after fitting the likelihood parameters by optimising the contribution of the reflection vector to the bound in Eq. 8 from the main paper

$$\hat{\boldsymbol{r}} = \arg\max_{\boldsymbol{r}:||\boldsymbol{r}||=1} \left\{ \sum_{j=1}^{W} \left( \boldsymbol{\psi}_{j'}^{d\top} (I - 2\boldsymbol{r}\boldsymbol{r}^\top) \bar{M}_j^d (I - 2\boldsymbol{r}\boldsymbol{r}^\top) \boldsymbol{\psi}_{j'}^d \right) \right\}. \tag{S.38}$$

This contribution has a similar form to those from the Wishart and log-Gaussian distributions, being fourth order in $\boldsymbol{r}$ with computable first and second derivatives, and can again be optimised using an interior point method.

#### S.2. DTI likelihood voxel volume correction

In the implementation details section of the main paper we highlight the fact that, although we resample the DTI images to the space of the structural imaging, the source dMRI is likely to have many fewer voxels. Hence, the process of upsampling can artifically inflate the contribution of the dMRI to the likelihood. In such cases it makes sense to re-weight the dMRI likelihood formulations to reflect the fact that the source dMRI is providing fewer measurements than the sMRI.

To do this we reformulate the objective function in Eq. 7 of the main paper, so that the diffusion likelihood is placed to the power of $\epsilon$,

$$\begin{aligned} O(\boldsymbol{\theta}|\boldsymbol{S},\boldsymbol{D},A,\boldsymbol{\gamma}) &= p(\boldsymbol{\theta}^a|\boldsymbol{\gamma}^a) + \sum_i^G \log p(\boldsymbol{\theta}_i^s|\boldsymbol{\gamma}_i^s) + \sum_j^W \log p(\boldsymbol{\theta}_j^d|\boldsymbol{\gamma}_j^d) \\ &+ \sum_v^V \log\left(\sum_c^C \left\{ p(l_v^c|A,\boldsymbol{\theta}^a)\left[\sum_i^G g_{c,i} p(\boldsymbol{s}_v|\boldsymbol{\theta}_i^s)\right]\left[\sum_j^W w_{c,j} p(\boldsymbol{d}_v|\boldsymbol{\theta}_j^d)\right]^\epsilon \right\}\right), \end{aligned} \tag{S.39}$$

where $\epsilon$ is the ratio between the volume of an sMRI voxel and a dMRI voxel.

Following the same derivation as in section 2.2 we can then find a lower bound for this objective function by applying Jensen's inequality twice, defining two separate soft segmentations at the current parameter estimates

$$\bar{q}_v^{c,i} = \frac{p(l_v^c|A,\boldsymbol{\theta}^a)p(\boldsymbol{s}_v|\boldsymbol{\theta}_i^s)\left[\sum_j w_{c,j} p(\boldsymbol{d}_v|\boldsymbol{\theta}_j^d)\right]^\epsilon}{\sum_c p(l_v^c|A,\boldsymbol{\theta}^a)\left[\sum_i g_{c,i} p(\boldsymbol{s}_v|\boldsymbol{\theta}_i^s)\right]\left[\sum_j w_{c,j} p(\boldsymbol{d}_v|\boldsymbol{\theta}_j^d)\right]^\epsilon} \qquad \sum_{c,i} g_{c,i}\bar{q}_v^{c,i} = 1,$$

$$\hat{q}_v^{c,j} = \frac{p(\boldsymbol{d}_v|\boldsymbol{\theta}_j^d)}{\sum_j w_{c,j} p(\boldsymbol{d}_v|\boldsymbol{\theta}_j^d)} \qquad \sum_j w_{c,j}\hat{q}_v^{c,j} = 1.$$

Substituting these in to Eq. (S.39) we obtain a new bound

$$\begin{aligned} O(\boldsymbol{\theta}|\boldsymbol{S},\boldsymbol{D},A,\boldsymbol{\gamma}) &\geq p(\boldsymbol{\theta}^a|\boldsymbol{\gamma}^a) + \sum_i^G \log p(\boldsymbol{\theta}_i^s|\boldsymbol{\gamma}_i^s) + \sum_j^W \log p(\boldsymbol{\theta}_j^d|\boldsymbol{\gamma}_j^d) + \sum_{v,c,i} g_{c,i}\bar{q}_v^{c,i}\left[\log\left[p(l_v^c|A,\boldsymbol{\theta}^a)p(\boldsymbol{s}_v|\boldsymbol{\theta}_i^s)\right]\right] \\ &+ \sum_{v,c,i,j} g_{c,i}\bar{q}_v^{c,i}\epsilon w_{c,j}\hat{q}_v^{c,j}\log p(\boldsymbol{d}_v|\boldsymbol{\theta}_j^d) - \sum_{v,c,i} g_{c,i}\bar{q}_v^{c,i}\left[\log\bar{q}_v^{c,i}\right] - \sum_{v,c,i,j} g_{c,i}\bar{q}_v^{c,i}\epsilon w_{c,j}\hat{q}_v^{c,j}\log\hat{q}_v^{c,j}, \end{aligned} \tag{S.40}$$

which can be further simplified by setting a single combined soft segmentation

$$\begin{aligned} q_v^{c,i,j} &= g_{c,i}w_{c,j}\bar{q}_v^{c,i}\hat{q}_v^{c,j} \\ &= \frac{g_{c,i}w_{c,j}p(l_v^c|A,\boldsymbol{\theta}^a)p(\boldsymbol{s}_v|\boldsymbol{\theta}_i^s)p(\boldsymbol{d}_v|\boldsymbol{\theta}_j^d)\left[\sum_j w_{c,j}p(\boldsymbol{d}_v|\boldsymbol{\theta}_j^d)\right]^{\epsilon-1}}{\sum_c p(l_v^c|A,\boldsymbol{\theta}^a)\left[\sum_i g_{c,i}p(\boldsymbol{s}_v|\boldsymbol{\theta}_i^s)\right]\left[\sum_j w_{c,j}p(\boldsymbol{d}_v|\boldsymbol{\theta}_j^d)\right]^\epsilon}, \end{aligned} \tag{S.41}$$

resulting in the new bound

$$\begin{aligned} O(\boldsymbol{\theta}|\boldsymbol{S},\boldsymbol{D},A,\boldsymbol{\gamma}) &\geq p(\boldsymbol{\theta}^a|\boldsymbol{\gamma}^a) + \sum_i^G \log p(\boldsymbol{\theta}_i^s|\boldsymbol{\gamma}_i^s) + \sum_j^W \log p(\boldsymbol{\theta}_j^d|\boldsymbol{\gamma}_j^d) + \sum_{v,c,i,j} q_v^{c,i,j}\left[\log\left[p(l_v^c|A,\boldsymbol{\theta}^a)p(\boldsymbol{s}_v|\boldsymbol{\theta}_i^s)p(\boldsymbol{d}_v|\boldsymbol{\theta}_j^d)^\epsilon\right]\right] \\ &- \sum_{v,c,i,j} q_v^{c,i,j}\left[\epsilon\log q_v^{c,i,j} - \epsilon\log w_{c,j} - \log g_{c,i}\right] - (1-\epsilon)\sum_{v,c,i,j} q_v^{c,i,j}\log\left[\sum_j q_v^{c,i,j}\right]. \end{aligned} \tag{S.42}$$

Extracting the contributions of structural and diffusion components $i$ and $j$ we then have the following equations to optimise for updated parameter estimates.

$$Q_s(\boldsymbol{\theta}_i^s) = \log p(\boldsymbol{\theta}_i^s|\boldsymbol{\gamma}_i^s) + \sum_v^V\left[\sum_{c,j} q_v^{c,i,j}\right]\log p(\boldsymbol{s}_v|\boldsymbol{\theta}_i^s), \tag{S.43}$$

$$Q_d(\boldsymbol{\theta}_j^d) = \log p(\boldsymbol{\theta}_j^d|\boldsymbol{\gamma}_j^d) + \epsilon\sum_v^V\left[\sum_{c,i} q_v^{c,i,j}\right]\log p(\boldsymbol{d}_v|\boldsymbol{\theta}_j^d). \tag{S.44}$$

#### S.3. Parameter initialisation

Here we expand on the details required for initialisation of the GEM algorithm. In general, during each E step we define a soft segmentation $q_v^{c,i,j}$ from our previous parameter estimates using the formula in Eq. (9). These are then summed across diffusion or structural components to provide component-wise soft segmentations $q_v^i$ for the structural components and $q_v^j$ for the diffusion components. We then use these component-wise soft segmentations to estimate the next set of parameters during the M step. However, at the start of the algorithm we need to initialise our first parameter estimates before generating the soft segmentation.

*Structural.* To perform this initialisation, we begin with the segmentation provided by Freesurfer's recon-all stream (*aseg.mgz*). We first initialise the deformation of the atlas by fitting our labels to this coarse automated segmentation. We then group labels according to their shared components in the structural model and erode the grouped labels, to account for errors in the aseg. These eroded labels are used to set the hyperparameters of the structural prior, using robust statistics, before setting our initial $q_v^i$ to 1 in the corresponding voxels and 0 elsewhere. We then perform structural only GEM for a number of iterations, using either the FA or log determinant of the DTI as an additional structural channel.

*log-Gaussian & DSW-beta.* After initialising the structural parameters we turn to the diffusion model. At this point, we first examine groups of label classes for which we specify a single diffusion component distribution. We generate an initial soft segmentation for this diffusion component by using the posteriors of the structural model

$$q_v^j = \frac{\sum_{c,i} g_{c,i} p(l_v^c|A, \boldsymbol{\theta}^a) p(\boldsymbol{s}_v|\boldsymbol{\theta}_i^s)}{\sum_{\{c,i\}} g_{c,i} p(l_v^c|A, \boldsymbol{\theta}^a) p(\boldsymbol{s}_v|\boldsymbol{\theta}_i^s)}. \tag{S.45}$$

However, for the classes in which there is a mixture of diffusion components, the process is more complex.

For the log-Gaussian and DSW-beta likelihood models we modify the soft segmentation in Eq. (S.45) by multiplying it with an initial, component-wise, segmentation. This gives

$$q_v^j = \frac{\sum_{c,i} g_{c,i} p(l_v^c|A, \boldsymbol{\theta}^a) p(\boldsymbol{s}_v|\boldsymbol{\theta}_i^s)}{\sum_{\{c,i\}} g_{c,i} p(l_v^c|A, \boldsymbol{\theta}^a) p(\boldsymbol{s}_v|\boldsymbol{\theta}_i^s)} \cdot \delta(\bar{l}_v = j) \tag{S.46}$$

where $\delta$ is the kronecker delta function and $\bar{l}_v$ is the initial component-wise segmentation. To generate $\bar{l}_V$ we apply k-means clustering to eroded label maps generated using Eq. (S.45). In the case of the log-Gaussian method we apply k-means clustering using the $L^1$, and for the DSW-beta model we use $1 - |\cos(\phi)|$, where $\phi$ is the angle between the two vectors being compared. In each case k-means is initialised from deterministic points picked using robust statistics.

*Wishart.* To initialise the Wishart likelihood model we use a different approach, instead of defining an initial component-wise segmentation using k-means we use techniques from Wishart mixture models of motion to retrieve initial parameter estimates. Following the work of Saint-Jean and Nielsen (2013, 2014) we use the k-MLE++ algorithm. We modify this k-MLE++ algorithm from the one proposed by Saint-Jean and Nielson, to ensure seed points are chosen in a deterministic way and to include our prior on the degrees of freedom parameter.

As a brief summary, the k-MLE++ algorithm makes use of the fact that the PDFs of exponential family distributions may be written in the form

$$p(x|\theta) = \exp\{\langle\theta, t(x)\rangle + k(x) - F(\theta)\}, \tag{S.47}$$

where $\theta$ holds some function of the distribution parameters, $\langle\cdot,\cdot\rangle$ is an inner product and $t(x)$ are the sufficient statistics for the distribution. In our case we set $\theta_n = {}^{(n-4)}/_2$, $\theta_V = V^{-1}$ and the sufficient statistics $t(\boldsymbol{d}) = (\log|\boldsymbol{d}|, -\frac{1}{2}\boldsymbol{d})$ giving the Wishart PDF, including gamma prior on degrees of freedom, as

$$p(\boldsymbol{d}|\theta_n, \theta_V, \alpha, \beta) = \exp\left\{\langle\theta_n, \log|\boldsymbol{d}|\rangle_{\mathbb{R}} + \langle\theta_V, -\tfrac{1}{2}\boldsymbol{d}\rangle_{HS} + k(\boldsymbol{d}, \alpha, \beta) - F(\theta_n, \theta_V, \alpha, \beta)\right\}, \tag{S.48}$$

with

$$F(\theta_n, \theta_V, \alpha, \beta) = (\theta_n + 2)(3\log(2) - \log|\theta_V|) + \log\Gamma_3(\theta_n + 2) - (\alpha - 1)\log(\theta_n + 1) + \beta(\theta_n + 1). \tag{S.49}$$

We can then define the Bregman divergence as

$$\begin{aligned}
B_{F^*}(p:q) &= F^*(p) - F^*(q) - \langle p-q, \nabla F^*(q)\rangle \\
F^*(q) &= \sup\{\langle q,\phi\rangle - F(\phi)\} \\
\nabla F^*(q) &= \arg\max_{\phi}\{\langle q,\phi\rangle - F(\phi)\} \\
&= \arg\max_{\phi}\{\langle q_n,\phi_n\rangle_{\mathbb{R}} + \langle q_V,\phi_V\rangle_{HS} - F(\phi_n,\phi_V)\}
\end{aligned} \tag{S.50}$$

So $\nabla F^*(t(\boldsymbol{d})) = \nabla F^*(\log|\boldsymbol{d}|, -\frac{1}{2}\boldsymbol{d})$ is the maximum likelihood estimate (*MLE*) for $\theta_n, \theta_V$ given the matrix $\boldsymbol{d}$ and $F^*$ is the log value of the PDF evaluated at this MLE excluding the contribution from $k(\boldsymbol{d})$. In this way we can use the Bregman divergence as a "distance" measure between the Wishart distributed tensors and seed points for each cluster. Combining Eq. (S.49) with the k-MLE++ algorithm from Saint-Jean and Nielsen (2014), and replacing random samplings with selections at the second and third quartiles, we define the deterministic k-MLE++ algorithm shown in Algorithm 1. Running this algorithm on eroded segmentations calculated using Eq. (S.45) results in initial parameter estimates for the Wishart distribution.

---

**Algorithm 1:** Deterministic k-MLE++

**Input :** A sample $\chi = \{\boldsymbol{d}_1, \ldots, \boldsymbol{d}_N\}$, $t(\boldsymbol{d}_i)$ the sufficient statistics, $F^*$ the dual log-normalizer of an exponential family, $K$ the number of clusters.

**Output:** Initial mixture MLE $\{\theta_n, \theta_v\}_{i=1}^K$

Choose first seed $\eta_1 = t(x_i)$ for $i = \arg\operatorname{median}_{i'} \log|\boldsymbol{d}_{i'}|$;

**for** $j = 2, \ldots, K+2$ **do**

  **foreach** $\boldsymbol{d}_i \in \chi$ **do**

    $p_i^j = B_{F^*}(t(\boldsymbol{d}_i) : \eta_{j-1})$;

    $p_i = \frac{\min_{j' \in [1,j-1]} p_i^{j'}}{\sum_{i'=1}^N \min_{j' \in [1,j-1]} p_{i'}^{j'}}$;

  **end**

  Choose $\eta_j \in \{t(\boldsymbol{d}_1), \ldots, t(\boldsymbol{d}_N)\}$ at third quartile of $p_i$;

**end**

**foreach** *combination* $\boldsymbol{\alpha}_k \in {}^{K+2}C_K$ *of* $\{\eta_j\}$ **do**

  $q_i^{\alpha_k} = \min_{j' \in \alpha_k} p_i^{j'}$;

  cost $\boldsymbol{\alpha}_k = \operatorname{median} q_i^{\alpha_k}$;

**end**

Choose $\boldsymbol{\alpha} = \arg\min_{\alpha_k} \operatorname{cost} \boldsymbol{\alpha}_k$;

Set $\{\theta_n, \theta_v\}_{i=1}^K = \{\nabla F^*(\eta_j)\}_{j \in \alpha}$

---

### S.4. Abbreviations

*S.4.1. General mathematical notation*

- $\boldsymbol{s}_v$ - voxel structural data
- $\boldsymbol{d}_v$ - voxel DTI data
- $\boldsymbol{\theta}^s$ - structural distribution parameters
- $\boldsymbol{\theta}^d$ - dMRI distribution parameters
- $A$ - atlas weights
- $\boldsymbol{\theta}^a$ - mesh position
- $l_v^c$ - label is class $c$ in voxel $v$
- $\boldsymbol{\gamma}$ - prior parameters for each distribution
- $g_{c,i}$ - mixture weight of structural component $i$ in class $c$ $(\sum_i g_{c,i} = 1)$
- $w_{c,j}$ - mixture weight of dMRI component $j$ in class $c$ $\left(\sum_j w_{c,i} = 1\right)$
- $G$ - number of structural distribution components
- $W$ - number of dMRI distribution components
- $V$ - number of voxels
- $c$ - class index
- $C$ - number of classes
- $\epsilon$ - ratio of voxel volumes between sMRI and dMRI ($\epsilon < 1$ when the dMRI is lower resolution than the sMRI)

#### *S.4.2. Thalamic labels*

Table S.1: Summary of the correspondences between the Grouped labels used for assessment, the manual label protocol and the histological labels output by our algorithm. We include the full name of each nucleus here for reference. For a full histological definition of each label please see Table 2 in Iglesias et al. (2018).

| Grouping | Manual label | Abbreviation | Nucleus |
|---|---|---|---|
| Antero-Lateral | Anterior | AV | Anteroventral |
| | Dorsal | LD | Laterodorsal |
| | | LP | Lateral posterior |
| | Lateral-Rostral | VA | Ventral anterior |
| | | VAmc | Ventral anterior magnocellular |
| | | VLa | Ventral lateral anterior |
| | | VLp | Ventral lateral posterior |
| | | VM | Ventromedial |
| Lateral-Caudal | Lateral-Caudal | VPL | Ventral posterolateral |
| Intralaminar | Intralaminar | CeM | Central medial |
| | | Pc | Paracentral |
| | | Pf | Parafascicular |
| | | MV(Re) | Reuniens (medial ventral) |
| | | Pt | Paratenial |
| | Intralaminar-Posterior | CM | Centromedian |
| Medial | Medial | CL | Central lateral |
| | | MDl | Mediodorsal medial magnocellular |
| | | MDm | Mediodorsal lateral parvocellular |
| Posterior | Pulvinar | L-Sg | Limitans (suprageniculate) |
| | | PuA | Pulvinar anterior |
| | | PuI | Pulvinar inferior |
| | | PuL | Pulvinar lateral |
| | | PuM | Pulvinar medial |
| | LGN | LGN | Lateral geniculate |
| | MGN | MGN | Medial Geniculate |

#### S.5. Expanded model comparison

In figure 6 from the main paper we displayed a comparison of segmentations from three methods on two subjects. The methods compared were the whole thalamus segmentation generated by Freesurfer's recon-all stream, a structural Bayesian segmentation and a joint segmentation generated using our DSW-beta likelihood model. Here we provide further comparison figures to give a more complete impression of the competing method quality on HCP. Figures S.1 and S.2 compare segmentations of subject 1 from figure 6 for five competing methods. These methods are:

1. structural only segmentation using the Freesurfer default implementation,
2. structural only segmentation with atlas mesh stiffness optimised for HCP as was used for the main paper results and figures,
3. joint structural and diffusion segmentation using the Wishart likelihood model,
4. joint structural and diffusion segmentation using the log-Gaussian likelihood model,
5. joint structural and diffusion segmentation using the DSW-beta likelihood model.

Additionally for the joint segmentation models we have included a division in the medial pulvinar to demonstrate the medial/lateral division of the PuM used in defining the component likelihood models.

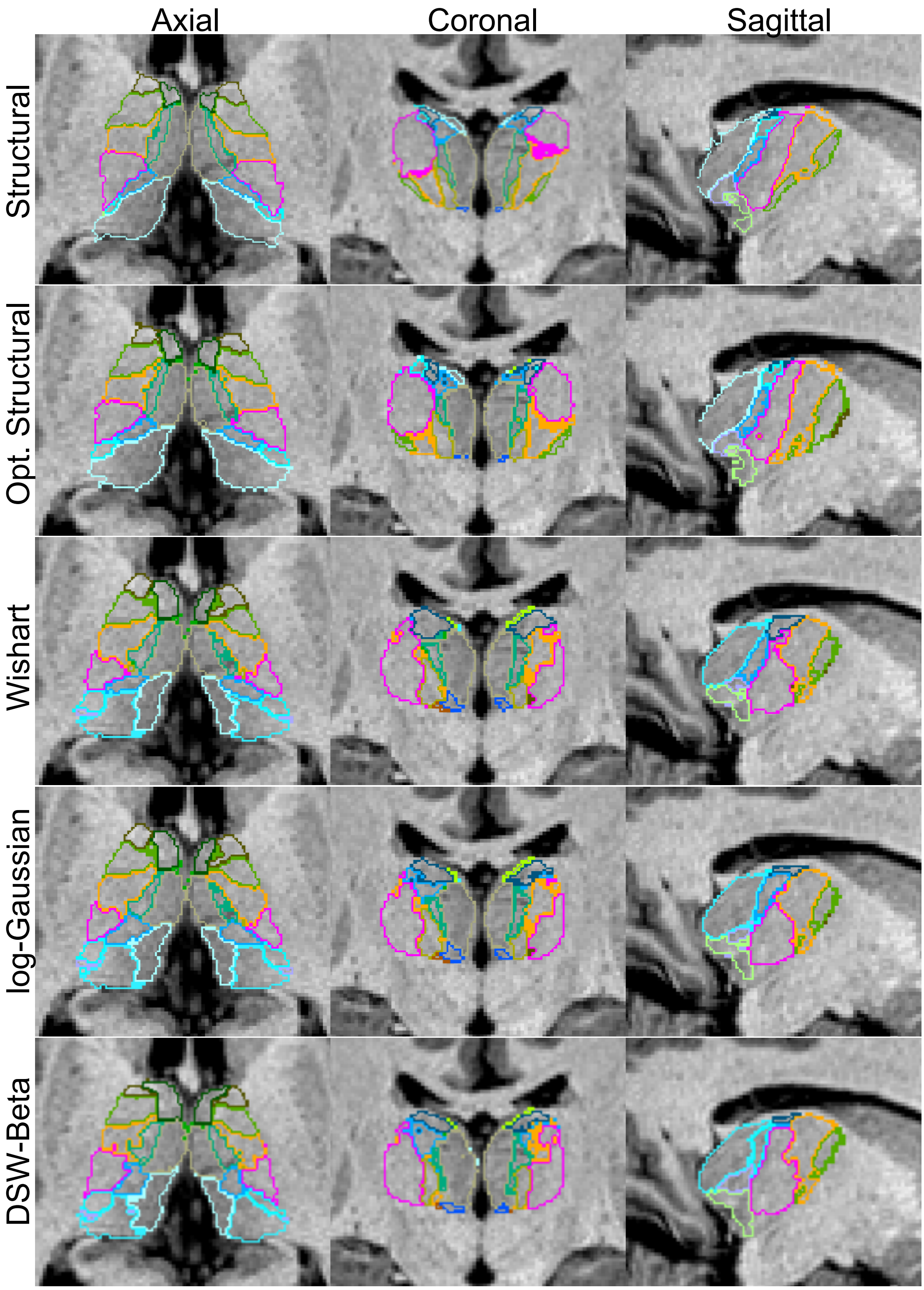


Figure S.1: Comparison of structural and joint segmentation of subject 1 from Figure 6 of the main paper. Structural, optimised structural, Wishart, log-Gaussian and DSW-beta models are compared, overlaid on T1 weighted structural MRI.

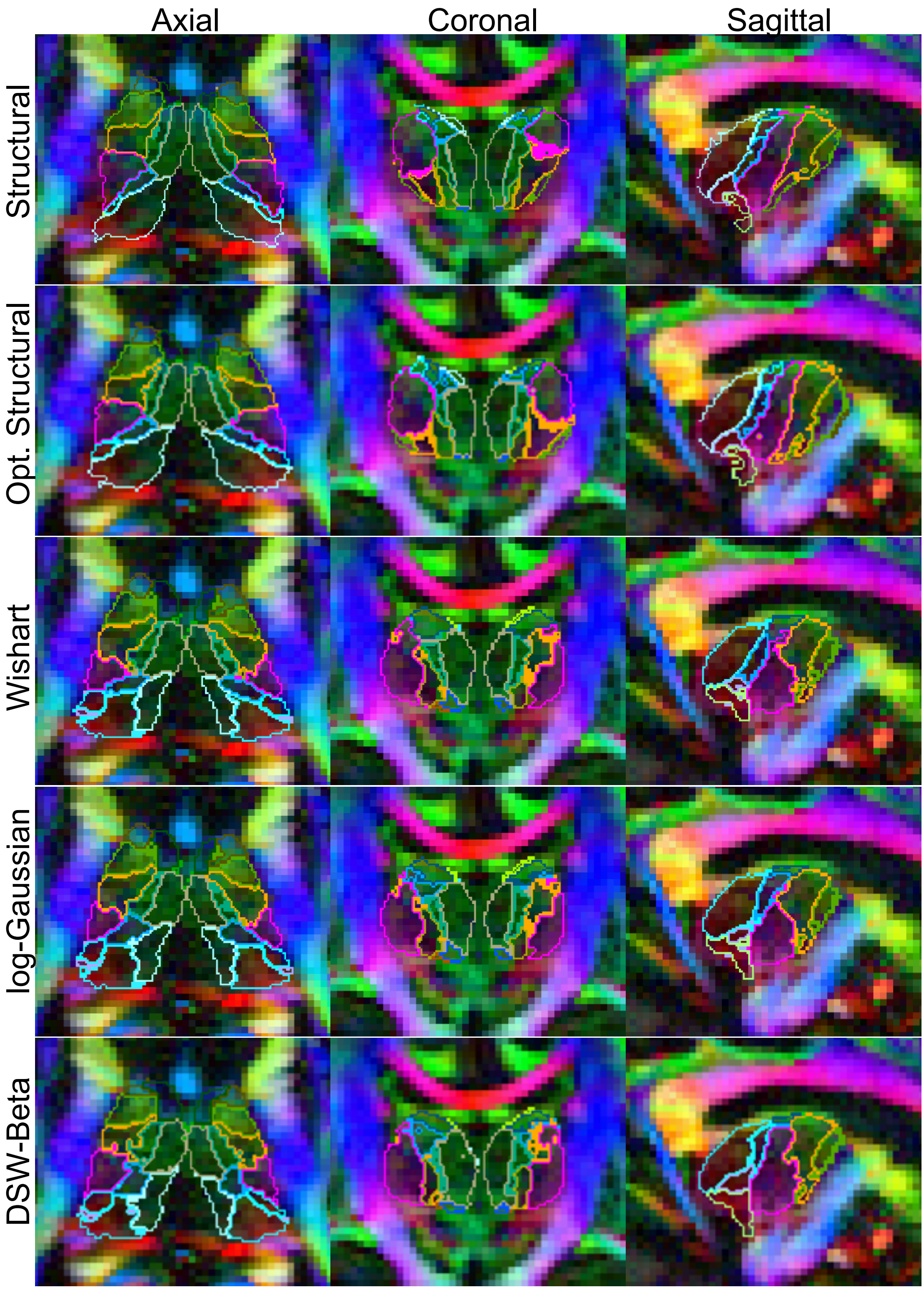


Figure S.2: Comparison of structural and joint segmentation of subject 1 from Figure 6 of the main paper. Structural, optimised structural, Wishart, log-Gaussian and DSW-beta models are compared, overlaid on DEC-FA.

#### Acknowledgments

This work was primarily funded by Alzheimer's Research UK (ARUK-IRG2019A003). The Dementia Research Centre is supported by Alzheimer's Research UK, Alzheimer's Society, Brain Research UK, and The Wolfson Foundation. This work was supported by the National Institute for Health Research (NIHR) Queen Square Dementia Biomedical Research Unit and the University College London Hospitals Biomedical Research Centre, the Leonard Wolfson Experimental Neurology Centre (LWENC) Clinical Research Facility, and the UK Dementia Research Institute, which receives its funding from UK DRI Ltd, funded by the UK Medical Research Council, Alzheimer's Society and Alzheimer's Research UK. MB is supported by a Fellowship award from the Alzheimer's Society, UK (AS-JF-19a-004-517). MB's work was also supported by the UK Dementia Research Institute which receives its funding from DRI Ltd, funded by the UK Medical Research Council, Alzheimer's Society and Alzheimer's Research UK. JDR is supported by the Miriam Marks Brain Research UK Senior Fellowship and has received funding from an MRC Clinician Scientist Fellowship (MR/M008525/1) and the NIHR Rare Disease Translational Research Collaboration (BRC149/NS/MH). JEI is supported by the European Research Council (Starting Grant 677697, project BUNGEE-TOOLS) and the NIH (1RF1MH123195-01 and 1R01AG070988-01).
